# Inherited DNA Repair Defects in Colorectal Cancer

**DOI:** 10.1101/256917

**Authors:** Saud H. AlDubayan, Marios Giannakis, Nathanael D. Moore, G. Celine Han, Brendan Reardon, Tsuyoshi Hamada, Xinmeng Jasmine Mu, Reiko Nishihara, Zhirong Qian, Li Liu, Matthew B. Yurgelun, Sapna Syngal, Levi A. Garraway, Shuji Ogino, Charles S. Fuchs, Eliezer M. Van Allen

**Affiliations:** Department of Medical Oncology, Dana-Farber Cancer Institute and Harvard Medical School, Boston, MA, 02215; The Broad Institute of MIT and Harvard, Cambridge, MA, 02142; Division of Genetics, Brigham and Women’s Hospital, Boston, MA, 02115; Department of Medicine, King Saud bin Abdulaziz University for Health Sciences, Riyadh, Saudi Arabia; Indiana University School of Medicine, Indianapolis, IN, 46202; Howard Hughes Medical Institute, Chevy Chase, MD, 20815; Program in MPE Molecular Pathological Epidemiology, Department of Pathology, Brigham and Women's Hospital, and Harvard Medical School, Boston, MA, 02115; Department of Oncologic Pathology, Dana-Farber Cancer Institute, Boston, MA, 02215; Department of Epidemiology, Harvard T.H. Chan School of Public Health, Boston, MA, 02115; Yale Cancer Center, Yale School of Medicine, New Haven, CT, 06510; Channing Division of Network Medicine, Department of Medicine, Brigham and Women's Hospital, Boston, MA, 02115

**Author notes:** Equal contribution (Saud H. AlDubayan, Marios Giannakis, Levi A. Garraway, Shuji Ogino, Charles S. Fuchs, Eliezer M. Van Allen).

**Keywords:** colorectal cancer genetics, germline genetics, cancer heritability, *ATM* mutations, *PALB2* mutations, homologous recombination, DNA repair deficiency

## Abstract

Colorectal cancer (CRC) heritability has been estimated to be around 30%. However, mutations in the known CRC susceptibility genes explain CRC risk in under 10% of the cases. Germline mutations in DNA-repair genes (DRGs) have recently been reported in CRC but their contribution to CRC risk is largely unknown. We evaluated the gene-level germline mutation enrichment of 40 DRGs in 680 unselected CRC individuals compared to 27728 ancestry-matched cancer-free adults. Significant findings were then examined in independent cohorts of 1661 unselected CRC cases and 1456 early-onset CRC cases. Of 680 individuals in the discovery set, 31 (4.56%) individuals harbored germline pathogenic mutations in known CRC susceptibility genes while another 33 (4.85%) individuals had DRG mutations that have not been previously associated with CRC risk. Germline pathogenic mutations in *ATM* and *PALB2* were enriched in both the discovery (OR= 2.81; P= 0.035 and OR= 4.91; P= 0.024, respectively) and validation sets (OR= 2.97; Adjusted P= 0.0013 and OR= 3.42; Adjusted P= 0.034, for *ATM* and *PALB2* respectively). Biallelic loss of *ATM* was evident in all cases with matched tumor profiling. CRC cases also had higher rates of actionable mutations in the HR pathway that can substantially increase the risk of developing cancers other than CRC. Our analysis provides evidence for *ATM* and *PALB2* as CRC risk genes, underscoring the importance of the homologous recombination pathway in CRC. In addition, we identified frequent complete homologous recombination deficiency in CRC tumors, representing a unique opportunity to explore targeted therapeutic interventions such as PARPi.

## Introduction

Colorectal cancer (CRC) [MIM: 114500] is the third most common malignancy in the US^1^. Although most CRC cases are thought to be sporadic, recent twin studies have estimated that 30% of the inter-individual variability in CRC risk is attributed to inherited genetic factors^2^. Over the past few decades, several CRC predisposition genes, including *APC* [MIM: 611731], *MLH1* [MIM: 120436], *MSH2* [MIM: 609309], *MSH6* [MIM: 600678], *PMS2* [MIM: 600259], *STK11* [MIM: 602216], *MUTYH* [MIM: 604933], *SMAD4* [MIM: 600993], *BMPR1A* [MIM: 601299], *PTEN* [MIM: 601728], *TP53* [MIM: 191170], *CHEK2* [MIM: 604373], *POLD1* [MIM: 174761] and *POLE* [MIM: 174762], have been described^3^^-^^5^. Collectively, mutations in these Mendelian CRC risk genes explain the increased risk for CRC in 5-10% of unselected cases^6^^-^^9^. The discrepancy between the proportion of CRC cases explained by these genetic risk factors and the estimated degree of heritability, known as “missing heritability”, indicates that one or more undiscovered inherited risk factors contribute to CRC risk.

DNA-repair is a critical biological process that prevents permanent DNA damage and ensures genomic stability. Although defects in DNA mismatch repair and certain DNA polymerases have been implicated in CRC risk, the role of other canonical DNA repair pathways is less defined. Our group and others have reported several observational studies which showed that some CRC cases were found to have germline mutations in DNA-repair genes (DRGs), such as *ATM* [MIM: 607585], *BRCA1* [MIM: 113705], *BRCA2* [MIM: 600185], and *PALB2* [MIM: 610355], that have classically been associated with susceptibility to cancers other than CRC^6, 10, 11^. As these DRG mutations are also present in the general population at a very low frequency, it is still unclear if these DRG defects are truly associated with a higher CRC risk or merely represent incidental findings in these CRC individuals^12^. To date, there has not been a case-control study to systematically examine candidate DRGs for potential germline mutation enrichment.

Here, we build upon our previous observations to evaluate the role of gene-level DRG defects in CRC susceptibility using germline data from CRC individuals and cancer-free controls in a case-cohort study, with complementary somatic analyses of candidate genes. We hypothesized that germline mutations in DRGs previously linked to other Mendelian forms of inherited cancer predisposition account for a significant fraction of the missing CRC heritability. To investigate this hypothesis, we studied germline whole exome sequencing data in a large discovery set of CRC cases who were not preselected for early-onset disease or positive family history and subsequently validated our findings in an independent large validation set of similarly unselected CRC cases. For CRC individuals who had disruptive germline mutations in genes related to homologous recombination, we also examined somatic tumor DNA for biallelic inactivation so as to explore whether such CRCs might theoretically be treated by agents that target deficient double-strand DNA repair (e.g. PARP inhibitors).

## Methods

### Study subjects

#### 1- Discovery set

Two independent cohorts that included 680 CRC persons were examined in the discovery phase (Figure S1). Of these, 591 CRC persons came from the population-based Nurses’ Health Study (NHS) and the Health Professionals Follow-up Study (HPFS) cohorts^13^. Only cases with available self-reported ancestry information were included in this case series. CRC cases from the NHS/HPFS were not selected on the basis of their age of presentation, stage of their disease or presence of a positive family history of CRC or other cancers^13^. In addition, 89 CRC persons from the CanSeq study at Dana-Farber Cancer Institute (DFCI) were included in the discovery set^14^. The CanSeq study is a single-arm prospective study that aims to evaluate the clinical utility of using paired (tumor and normal) whole exome sequencing in the clinical care of individuals with advanced cancer without pre-selection for early age at diagnosis or high-risk family histories (hereafter referred to as “unselected cases”)^15^. Both studies were approved by the Partners Human Research Committee institutional review board (NHS/HPFS: BWH IRB#2001-P-001945, CanSeq: DFCI IRB#12-078)), and informed consent was obtained from all subjects.

#### 2- Validation set

Germline data of 1661 subjects from two independent cohorts of unselected CRC cases, The Cancer Genome Atlas (TCGA; n = 603) and the cohort reported by Yurgelun et al. (n = 1058) were used to validate the main findings detected in the discovery phase (hereafter called “the validation set”)^16, 17^. Both cohorts were not selected for early-onset disease or positive family history. Similar variant calling and pathogenicity assessment pipelines were used to evaluate germline variants in both cohorts.

#### 3- Early-onset CRC set

To further delineate the penetrance of DRGs with significant germline mutation enrichment in the discovery and validation sets in CRC individuals, germline mutation enrichment in 1456 early-onset (age<56) CRC cases was evaluated. These cases were part of two large CRC studies^10, 18^. In total, our study evaluated relevant germline sequencing data of 3797 CRC cases relative to cancer-free adult controls (Figure S1).

### Sequencing and Bioinformatics Analysis

Germline DNA from the CRC subjects in the discovery set was obtained from whole blood or adjacent normal colon tissue that was dissected after pathology review. DNA was extracted from formalin-fixed, paraffin embedded (FFPE) blocks using commonly used practices^19^. All germline variants in the validation and early-onset CRC sets were detected form whole blood. Production pipelines of the germline variants of these cohorts are described in Table S1 and elsewhere^10, 13, 16-18^. Partial or whole gene deletions were not evaluated in this study.

### Selection of DNA-repair genes and gene sets

Only genes that have been clearly associated with a Mendelian cancer-predisposition syndrome in humans were examined. A total of 14 well-known CRC risk genes, as well as 40 DRGs that have been associated with cancer phenotypes other than CRC, were evaluated (Tables S2 and S3). Some of these DRGs such as *BLM* [MIM: 210900] and *NTHL1* [MIM: 602656] have been recently linked to CRC susceptibility, however these observations have not been so far independently validated so these genes were included in the DRG set to be evaluated here. Analysis of the germline variants in *POLE* and *POLD1* was restricted to the known pathogenic missense mutations in the exonuclease domain of the protein.

Of the examined DRGs, 14 genes play an important part in the homologous recombination pathway: *ATM*, *BARD1* [MIM: 601593], *BLM*, *BRCA1*, *BRCA2*, *BRIP1* [MIM: 605882], *MRE11* [MIM: 600814], *NBN* [MIM: 602667], *PALB2*, *RAD51* [MIM: 179617], *RAD51C* [MIM: 602774], *RAD51D* [MIM: 602954], *RAD54L* [MIM: 603615], and *XRCC3* [MIM: 600675]^20^. “Actionable DRGs” were defined as established cancer predisposition genes that confer a 3-fold or higher increase in the risk for cancer phenotypes other that CRC and for which enhanced screening and family genetic testing are recommended. Out of the examined DRGs, *ATM*, *BRCA1*, *BRCA2*, *BRIP1*, *PALB2*, *RAD51C*, and *RAD51D* were considered clinically actionable^21^^-^^25^.

### Variant Interpretation

An identical workflow for variant inclusion and pathogenicity assessment was used to evaluate the germline variants in both cases and controls (Table S1). The clinically-oriented American College of Medical Genetics and Genomics (ACMG) germline variant assessment guidelines were used to evaluate germline variants in cases and controls. Based on the available evidence, germline variants were classified into 5 categories: benign, likely benign, variants of unknown significance, likely pathogenic and pathogenic^26^. Only germline variants which had sufficient evidence of pathogenicity to be classified as pathogenic or likely pathogenic variants (hereafter collectively referred to as pathogenic mutations) were included. All variants of unknown significance (VUS) were excluded from all analyses.

### Frequency of mutations in the general population

Annotated germline variants in the examined genes in 53105 cancer-free adults from the Exome Aggregation Consortium (ExAC) (release 0.3.1 on 3/16/2016), excluding the TCGA cohort, were also evaluated using an identical workflow to the one used for cases ^27^. Frequencies of germline pathogenic mutations in the genes of interest were calculated for each of the continental populations in ExAC. Gene mutation frequencies for the ExAC Non-Finnish European (n=27173) and African & African American (n=4533) cohorts were then used to calculate the predicted pathogenic gene mutation frequency in an ancestry-matched control cohort of 27728 individuals (98%; 27173 Non-Finnish Europeans (NFE), and 2%; 555 African Americans (AFR)) (Figure S2)^28^. Population-specific common variant frequencies were similar in cases and controls decreasing the likelihood of a significant population structure (Figure S3). Ancestry information for some individuals in the validation set was not readily available. Since the majority of the cases included in these studies are expected to have European ancestry, non-Finnish European individuals from the ExAC cohort (ExAC_NFE; n= 27173) were used as a control group.

### Tumor LOH analysis

MuTect was applied to identify somatic single-nucleotide variants (SNVs)^29^. Strelka was used to detect small insertions and deletions. Individual sites were reviewed with Integrated Genomics Viewer (IGV)^30^. Using filtered-based method, artifacts from DNA oxidation during sequencing were removed^31, 32^. Annotation of identified variants was performed using Oncotator^33^. Probability distributions of possible cancer cell fractions (CCFs) of mutations were calculated, based on local copy-number and the estimated sample purity, using ABSOLUTE^34^.

### Statistical Analysis

A logistic regression model was used to examine the clinical characteristics of CRC cases with germline pathogenic mutations. Two-sided Fisher’s exact tests were used to calculate the odds ratios and confidence intervals (using “Minimum likelihood correction”) for the enrichment of germline pathogenic mutations in each of the examined DRGs. In addition, Exact binomial test of proportions was used to calculate the P value for the measured enrichment of each gene in CRC cases compared with the reference population. Consistent with established statistical methods for two-stage association studies, we implemented a permissive first discovery stage analysis where genes with P values smaller than 0.05 were considered significant. These top candidate genes were then tested in a subsequent validation phase in an independent cohort, prior to performing secondary analyses, with appropriate correction for multiple testing using Bonferroni correction^35^^-^^37^.

## Results

### Cohort characteristics and sequencing metrics of CRC cohorts

Demographic characteristics of all 680 CRC cases from the discovery cohort are summarized in Tables 1 and S4. The average target coverage for germline WES for the discovery set was 71.69X (NHS/HPFS) and 137.11X (CanSeq). DNA-repair genes, where significant germline pathogenic mutation enrichment was seen in the discovery set, were subsequently examined in 1661 unselected CRC cases and 1456 early-onset CRC cases (methods) ^10, 16^. Examined DRGs had an average coverage of 58.67X in the ExAC cohort (Figure S4 and Table S5).

**Table 1:**
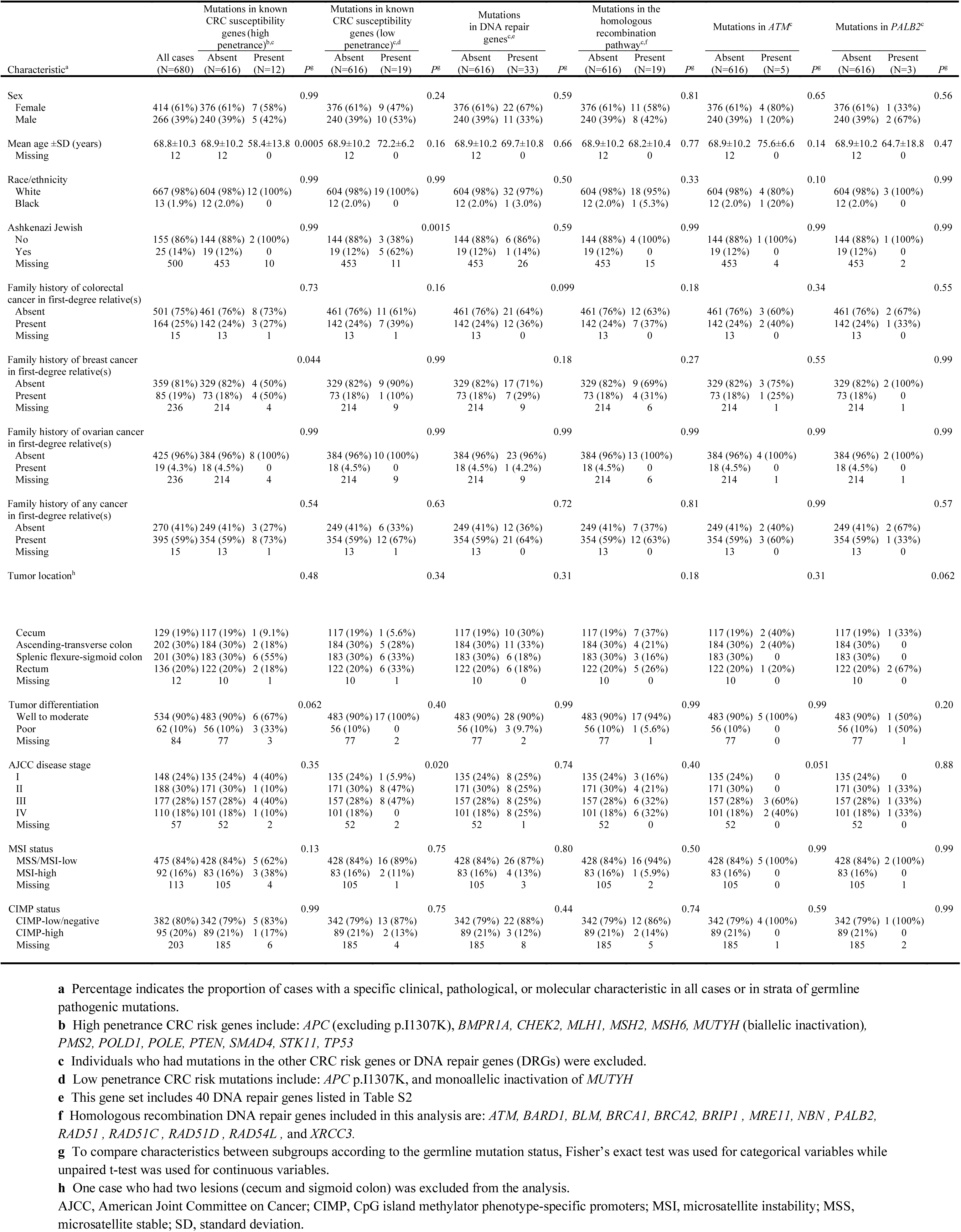
Clinical, pathological, and molecular characteristics of 680 colorectal cancer cases who were examined in the discovery set.

### Germline pathogenic mutations in known CRC risk genes

In the discovery set (n = 680), 31 (4.56%) individuals had germline CRC risk mutations. Of these, 12 (1.76%) harbored highly or moderately penetrant germline pathogenic mutations in *APC* (n=2), *CHEK2* (n=4), *MSH2* (n=1), *MSH6* (n=1), *PMS2* (n=2), and *TP53* (n=2) (Figures 1a and S5; Table S6). In addition, 19 (2.79%) individuals carried heterozygous germline pathogenic mutations in *MUTYH* (n=11, 1.62%) or the Ashkenazi founder low-penetrance variant, p.Ile1307Lys, in *APC* (n=8, 1.18%). Of 1661 unselected CRC individuals in the validation set, 93 (5.6%) individuals had at least one germline mutation in the CRC susceptibility genes (Figure 1a; Tables S7 and S8). The frequency of germline mutations in the mismatch repair genes (*MLH1*, *MSH2*, *MSH6* and *PMS2*) in the discovery CRC set (4 patients; 0.6%) is considerably lower than the frequency of these gene mutations in other studies^38^. This underrepresentation of Lynch syndrome patients in our discovery cohort could be attributed to the population-based nature of the NHS/HPFS cohorts as well as to the fact that these studies only enrolled cancer-free subjects, sometimes at a more advanced age for some individuals.

**Figure 1:**
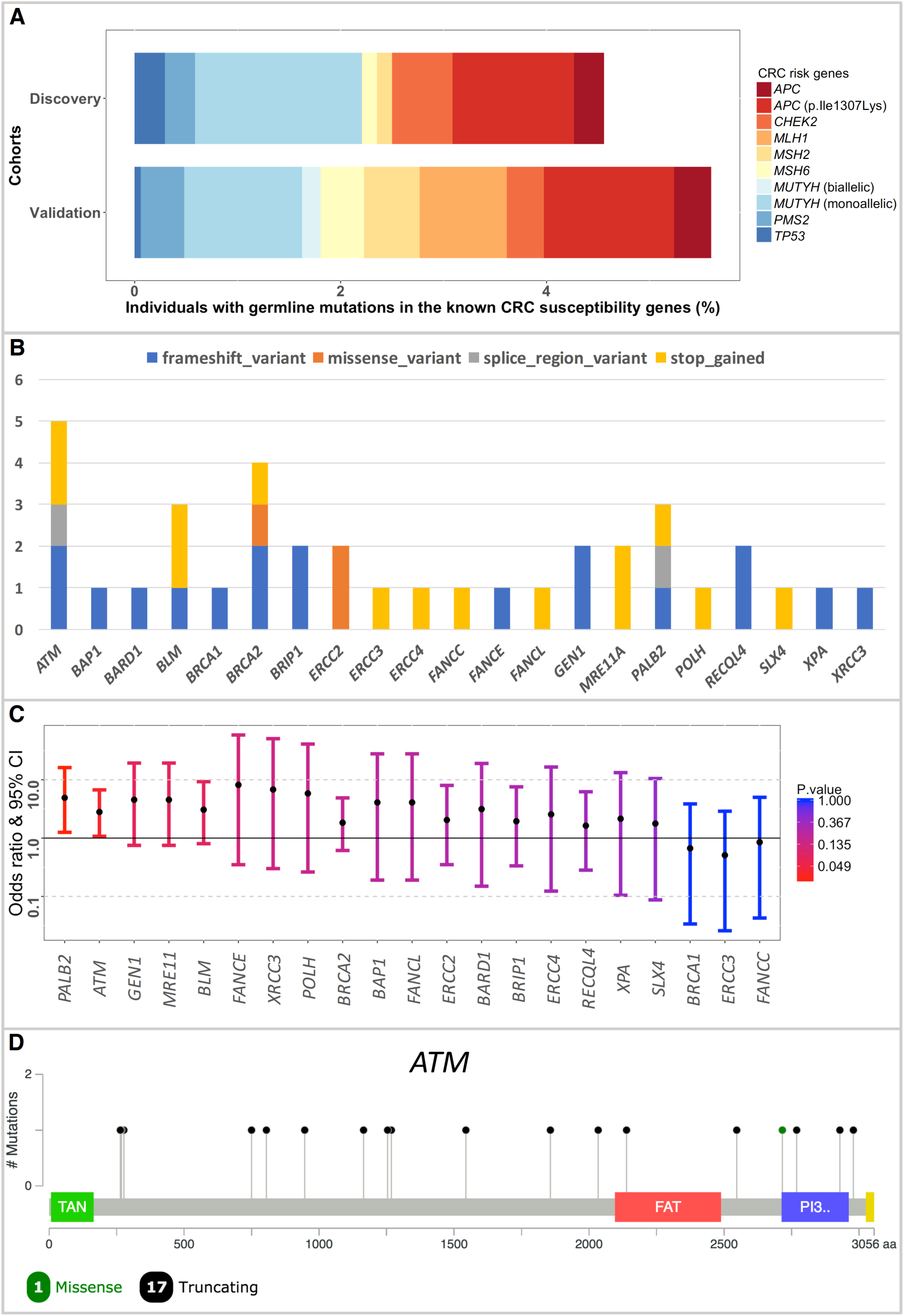
Germline pathogenic mutations in the known CRC predisposition genes and additional DNA repair genes. A; Proportions of cases with germline pathogenic mutations in the CRC risk genes in 680 CRC individuals in the discovery set and 1661 CRC cases in the validation set. B; Number and class of the detected germline pathogenic mutations in the DRGs in the discovery set (n=680). DRGs where no mutations were detected (n=19) are not shown here. C; Enrichment of germline pathogenic DRGs mutations in 680 CRC individuals in the discovery set. Fisher’s exact test was used to calculate the ORs and 95% confidence intervals. Two-sided binomial test was used to calculate the P values. D; A total of 18 germline pathogenic *ATM* mutations were seen in the discovery and validation sets in our study. This includes seven (38.9%) nonsense mutations, six (33.3%) frameshift mutations, three (16.6%) splice-site mutations, one (5.6%) known pathogenic in-frame deletion and one (5.6%) known pathogenic missense mutation.

### Germline pathogenic mutations in additional DNA-Repair genes

Next, germline variants in 40 DRGs in the discovery CRC set (n=680) were evaluated for pathogenicity. Thirty-three (4.85%) subjects had at least one germline pathogenic mutation in 21 of these DRGs (Figure 1b). Four (0.59%) individuals had 2 germline pathogenic mutations each in different DRGs (Table S9). There were no cases with germline pathogenic mutations in both sets of known CRC risk genes and the additional DRGs. Enrichment analysis of the discovery CRC set, relative to cancer-free individuals, showed significant germline pathogenic mutation enrichment in *ATM* and *PALB2* (Figure 1c; Table 2).

**Table 2:** Enrichment of germline pathogenic mutations in 680 CRC cases (discovery set) relative to 27728 ancestry-matched cancer-free adults from the ExAC cohort. Only genes with detected germline pathogenic mutations in cases are shown. (ExAC: Exome Aggregation Consortium)

| Gene | Cases with mutations in the discovery cohort (n= 680) | Prevalence of cases with mutations in the discovery cohort (%) | Cases with mutations in ancestry-matched control group (n=27728) | Prevalence of mutations in the control group (%) | Enrichment of pathogenic mutations in the discovery cohort (OR; Fisher's Exact test) | 95% Confidence Intervals (Fisher's Exact test) | P value (two-sided Exact Binomial test) |
| --- | --- | --- | --- | --- | --- | --- | --- |
| *ATM* | 5 | 0.74% | 73 | 0.26% | 2.81 | 1.07-6.71 | 0.035 |
| *BAP1* | 1 | 0.15% | 10 | 0.04% | 4.08 | 0.19-27.85 | 0.218 |
| *BARD1* | 1 | 0.15% | 13 | 0.05% | 3.14 | 0.15-19.04 | 0.273 |
| *BLM* | 3 | 0.44% | 40 | 0.14% | 3.07 | 0.8-9.28 | 0.077 |
| *BRCA1* | 1 | 0.15% | 61 | 0.22% | 0.67 | 0.03-3.86 | 1 |
| *BRCA2* | 4 | 0.59% | 89 | 0.32% | 1.84 | 0.61-4.89 | 0.177 |
| *BRIP1* | 2 | 0.29% | 42 | 0.15% | 1.94 | 0.33-7.57 | 0.275 |
| *ERCC2* | 2 | 0.29% | 40 | 0.14% | 2.04 | 0.35-8.00 | 0.25 |
| *ERCC3* | 1 | 0.15% | 80 | 0.29% | 0.51 | 0.03-2.89 | 1 |
| *ERCC4* | 1 | 0.15% | 16 | 0.06% | 2.55 | 0.12-16.55 | 0.325 |
| *FANCC* | 1 | 0.15% | 48 | 0.17% | 0.85 | 0.04-5.0 | 1 |
| *FANCE* | 1 | 0.15% | 5 | 0.02% | 8.16 | 0.35-58.6 | 0.115 |
| *FANCL* | 1 | 0.15% | 10 | 0.04% | 4.08 | 0.19-27.85 | 0.218 |
| *GEN1* | 2 | 0.29% | 18 | 0.06% | 4.54 | 0.75-19.35 | 0.073 |
| *MRE11* | 2 | 0.29% | 18 | 0.06% | 4.54 | 0.75-19.35 | 0.073 |
| *PALB2* | 3 | 0.44% | 25 | 0.09% | 4.91 | 1.26-16.19 | 0.024 |
| *POLH* | 1 | 0.15% | 7 | 0.03% | 5.83 | 0.26-40.98 | 0.158 |
| *RECQL4* | 2 | 0.29% | 50 | 0.18% | 1.63 | 0.28-6.23 | 0.347 |
| *SLX4* | 1 | 0.15% | 23 | 0.08% | 1.77 | 0.09-10.49 | 0.431 |
| *XPA* | 1 | 0.15% | 19 | 0.07% | 2.15 | 0.1-13.26 | 0.373 |
| *XRCC3* | 1 | 0.15% | 6 | 0.02% | 6.8 | 0.3-50.67 | 0.137 |

### Germline pathogenic mutations in *ATM*

Among 680 unselected CRC individuals, five (0.74%) had mutations in *ATM*. Germline mutations in *ATM* were significantly more prevalent in the CRC discovery set than cancer-free individuals (OR= 2.81; 95% CI= 1.07-6.71; P= 0.035) (Table S9). The frequency of *ATM* germline pathogenic mutations in the CanSeq cohort was not significantly higher than that of the NHS/HPFS cohort (P= 0.5) (Figure S6). Analysis of *ATM* mutation frequency in another 1661 unselected CRC cases, from the validation set, also identified significant enrichment of *ATM* germline pathogenic mutations (13 cases; 0.78%; OR= 2.97; 95% CI= 1.57-5.39; Adjusted P= 0.0013) (Figures 1d and 2a; Tables S10 and S11). Evaluation of an independent cohort of 1456 early-onset CRC individuals similarly showed significant enrichment of germline *ATM* mutations in these individuals (10 cases; 0.69%; OR= 2.6; 95% CI= 1.3-5.07; Adjusted P= 0.013) (Figure 2a).

**Figure 2:**
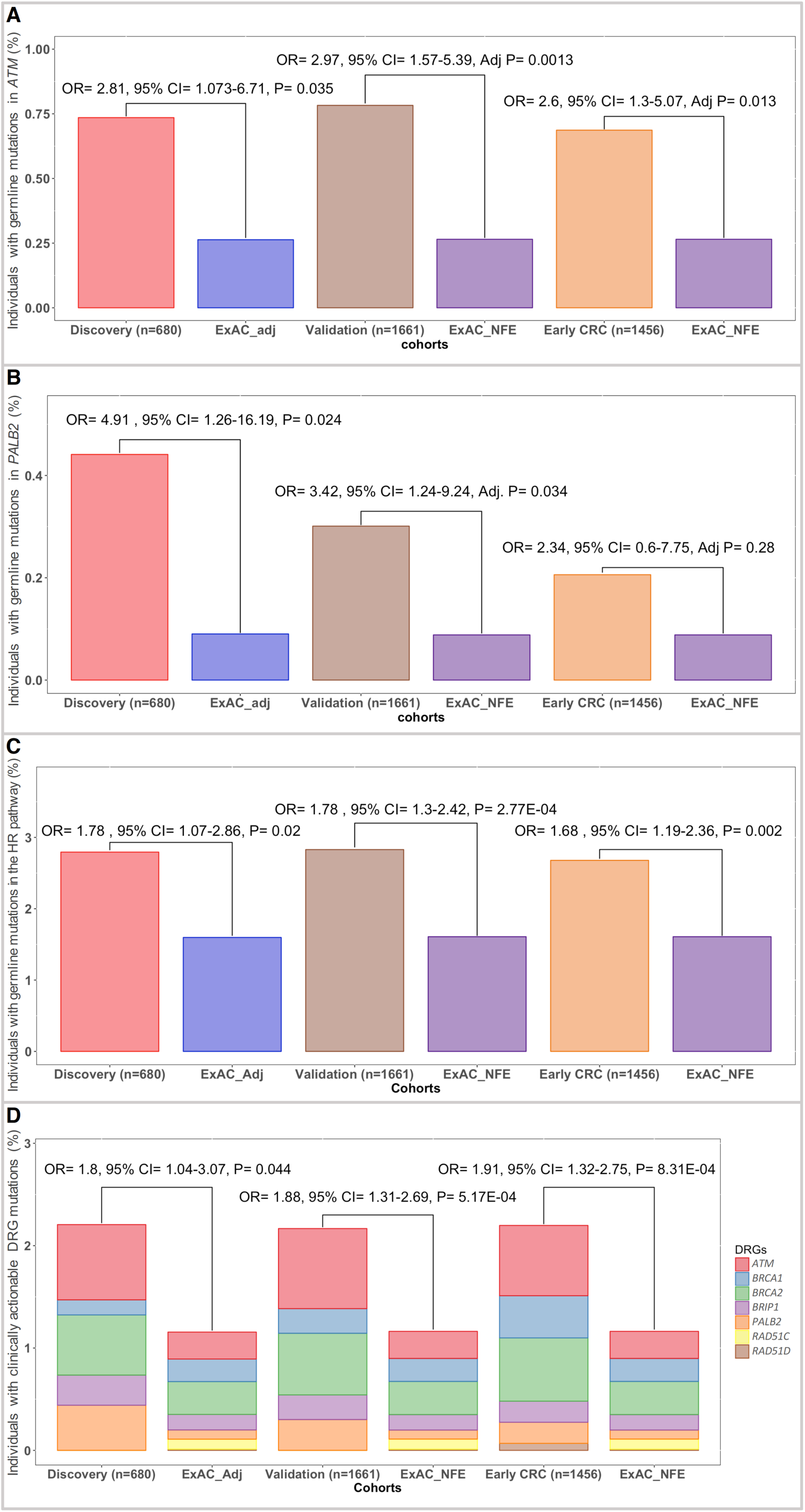
Enrichment of DRG mutations in various cohorts. A; Inherited pathogenic germline mutations in *ATM* were more commonly seen in individuals with CRC in the discovery, validation and early-onset CRC sets (n=680; n=1661, n=1456, respectively) compared with cancer-free individuals. B; Germline pathogenic mutations in *PALB2* were significantly enriched in unselected CRC cases from the discovery and validation sets. However, no significant enrichment was seen in the early-onset CRC cases. C; A secondary analysis of the homologous recombination pathway showed significant enrichment of germline HR gene mutations, as an aggregate, in all CRC cohorts. D; Individuals with CRC were also almost twice more likely to carry a clinically actionable mutation where screening recommendation do exist and which can greatly impact the clinical care offered to these individuals and their families.

Although most of the cases included in our study were of European ancestry, self-reported ancestry information, as previously shown, can be inaccurate^39^. To evaluate for spurious *ATM* mutation enrichment that could have resulted from inadequate population stratification, we next blinded the ancestry data of the CRC subjects from the validation cohort and examined *ATM* mutation enrichment relative to cancer-free controls from various continental populations in ExAC. Our analysis showed that regardless of the selected control population, rates of germline *ATM* mutations were significantly higher in the CRC validation set (n=1661) (OR= 2.4-6.5, Adjusted P< 0.05 for all pairwise comparisons; Binomial Exact with Bonferroni correction for 6 independent tests) (Figure S7).

### Germline pathogenic mutations in *PALB2*

Three individuals in our discovery cohort were found to have germline *PALB2* mutations, which represented a significant enrichment, compared to cancer-free controls (0.44%; OR= 4.91; 95% CI= 1.26-16.19; P= 0.024) (Table S9). This enrichment was also evident in 1661 unselected CRC cases from the validation cohort (5 cases; 0.3%; OR= 3.42; 95% CI= 1.24-9.24; Adjusted P= 0.034) (Figure 2b and Tables S10 and S11). Interestingly, no significant enrichment of germline *PALB2* mutations was seen in 1456 early-onset CRC cases (3 cases; 0.2%; OR 2.34; 95% CI= 0.6-7.75; Adjusted P= 0.28), suggesting late-onset penetrance of *PALB2* mutations in CRC individuals.

### Somatic loss of heterozygosity (LOH)

Matched tumor WES for most of the individuals with germline mutations in the discovery set (n=64) were available and examined for somatic loss of heterozygosity (LOH) (Table S12). Among the CRC risk genes, somatic inactivation of the wild-type allele was seen in *APC* (8 cases; 80%), *CHEK2* (1 case; 25%), *ERCC2* (2 case, 100%), *MSH2* (1 case; 100%), *MSH6* (1 case; 100%), *MUTYH* (2 cases; 18%), *PMS2* (2 case; 100%) and *TP53* (2 cases; 100%). Out of the examined DRGs, all individuals with germline pathogenic mutations in *ATM* (5; 100%) had evidence of somatic inactivation of the wild-type allele in the matched tumor samples (Figure S8). Somatic inactivation of the *ATM* wild-type allele, in all tumors with germline *ATM* events, provides compelling evidence for *ATM* to be etiologic for the development of CRC in these cases. No somatic LOH was detected in any of the tumors of individuals with germline *PALB2* mutations, though disruptive non-coding genetic and epigenetic events are not captured by tumor WES.

### Germline pathogenic mutations in the homologous recombination (HR) pathway

Given the observed mutations specifically in HR genes (*ATM* and *PALB2*), we next examined the frequency of inherited mutations affecting any of HR cancer-predisposition genes (methods). Unselected CRC individuals in the discovery set had a higher rate of germline pathogenic mutations in the HR genes compared with cancer-free individuals (19 cases; 2.8%; OR= 1.77; 95% CI= 1.07-2.84; P= 0.02) (Table S9). Evaluation of the validation and early-onset CRC sets also showed that CRC cases were more likely to have inherited HR mutations (validation set: 47 cases; 2.8%; OR= 1.78; 95% CI= 1.30-2.43; P= 2.77E-04; early-onset set: 39 cases; 2.68%; OR= 1.68; 95% CI= 1.19-2.35; P= 0.002) (Figure 2c; Tables S10 and S11). This effect did not seem to be purely driven by *ATM* and *PALB2* mutations, as when excluded, there was a trend, that did not reach statistical significance, for germline disruptive events in other HR genes to be more prevalent in the CRC validation set compared with cancer-free adults (OR= 1.4; 95% CI= 0.95-2.06; P= 0.077) (Figure S9).

### Clinical actionability and risk of other cancers in CRC individuals

Analysis of mutations in actionable DRGs (*ATM*, *BRCA1*, *BRCA2*, *BRIP1*, *PALB2*, *RAD51C*, and *RAD51D*) in the discovery set identified a total of 15 germline pathogenic mutations in 14 (2.1%) CRC persons. One person had two actionable mutations in *BRCA2* and *PALB2*. Compared with cancer-free individuals, actionable cancer-risk mutations were approximately twice more prevalent in CRC cases from the discovery set (OR= 1.8; 95% CI= 1.04-3.07; P= 0.04), the validation set (36 cases; 2.17%; OR= 1.88; 95% CI= 1.31-2.69; P= 5.17E-04) as well as the early-onset CRC set (32 cases; 2.2%; OR= 1.91; 95% CI= 1.32-2.75; P= 8.31E-04) (Figure 2d).

### Utility of testing relevant DRGs in CRC

Collectively, CRC heritability in up to about 1.2% of unselected CRC cases may be explained by higher rates of mutations in *ATM* and *PALB2*. To examine the potential impact of performing germline testing of *ATM* and *PALB2* on diagnostic yield, we next examined the CRC-specific germline panels offered by eight of the largest commercial laboratories in the US (as of September 2017). In addition to the known CRC risk genes, our evaluation showed that germline analysis of *ATM* is only occasionally included in these panels whereas *PALB2* and other actionable DRGs are not captured by these clinical tests (Figure S10).

### Clinical characteristic of mutation carriers in the discovery set

Overall, there were no significant differences in clinical characteristics between DRG mutant or non-mutant CRC cases (Table 1). Although on average, CRC individuals with high penetrance germline CRC risk mutations presented 10.5 years younger that mutation-negative individuals (P= 0.0005), CRC individuals with germline pathogenic mutations in *ATM*, *PALB2*, the HR genes or DRGs were not more likely to present earlier that mutation-negative persons. All five germline *ATM* mutation carriers presented with stage III or IV disease (compared with 46% of mutation-negative CRC cases; P= 0.051) (Figure 3). Individuals with germline pathogenic mutations in CRC risk genes, the DRGs, *ATM* or *PALB2* were not more likely to report a first-degree family member with CRC or other cancer types (Figure S11). Interestingly, individuals carrying a high penetrance CRC risk mutations were more likely to report a positive family history of breast cancer.

**Figure 3:**
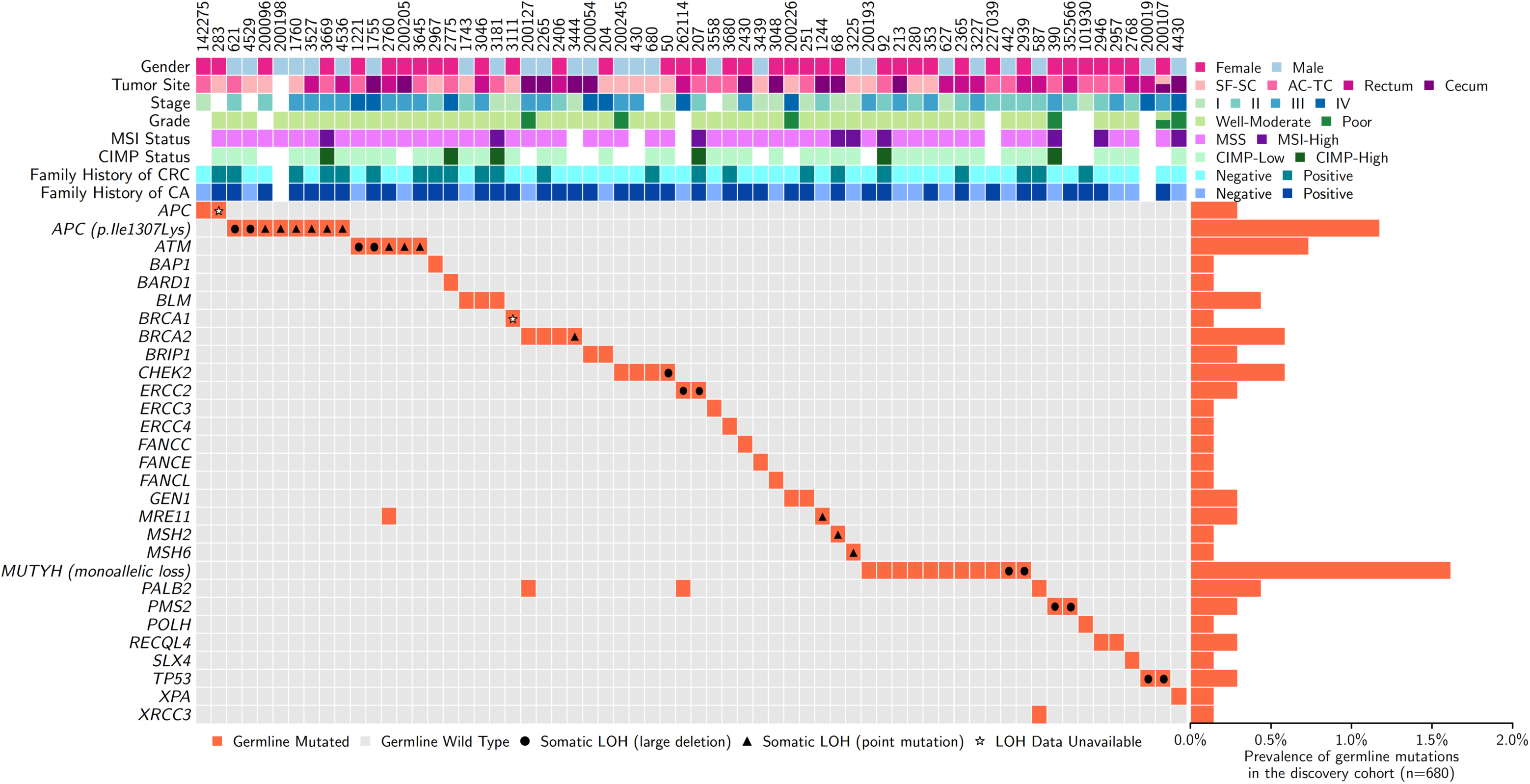
Clinical and molecular characteristics of all cases with germline pathogenic mutations in CRC risk genes and DRGs in our discovery set. All individuals with germline pathogenic mutations in *ATM* had somatic LOH in their tumor samples. Two of these cases had large deletions that affected the wild-type *ATM* allele while three had truncating point mutations leading to the loss of *ATM* wild-type allele as well. (AC-TC: ascending colon to transverse colon; SF-SC: splenic flexure to sigmoid colon; MSI: microsatellite instability; MSS: microsatellite stable; CIMP: CpG island methylator phenotype-specific promoters; LOH: loss of heterozygosity).

## Discussion

Most of the colorectal cancer heritability is still incompletely characterized. Mutations of several cancer-predisposition DRGs that are not typically associated with CRC have been recently reported in individuals with CRC, however, the clinical significance of these results has not been firmly established. Here, we present a systematic analysis of DRG mutations in large independent CRC cohorts relative to cancer-free adults to evaluate novel observations in known CRC susceptibility genes and to identify new CRC susceptibility genes.

We found that a gene-level analysis of DRGs revealed significantly higher rates of *ATM* mutations in CRC cases compared with cancer-free controls, going beyond observational studies to implicate its role as a novel CRC susceptibility gene. *ATM* is a master regulating kinase that is activated in response to DNA damage. Heterozygous carriers of *ATM* mutations have been reported to have a higher risk of breast [MIM: 114480] and potentially pancreatic cancer [MIM: 260350]^11^. A previous cohort-based study that evaluated the risk of various cancers in families of individuals with ataxia telangiectasia [MIM: 208900], which results from biallelic loss of *ATM*, showed no increased risk of CRC in the obligate carrier parents of these cases. However, a secondary analysis in that study showed that, collectively, there was an increased risk of CRC when all the heterozygous *ATM* carrier relatives were evaluated (RR=2.54, 95% CI= 1.06-6.09), though this association was not statistically significant once corrected for multiple hypothesis testing^11^. A larger subsequent study on *ATM* carriers also failed to detect any enrichment of CRC events in heterozygous *ATM* carries^40^. However, a recent GWAS that evaluated three loss-of-function *ATM* variants in several cancer phenotypes showed a higher risk for CRC in cases (OR=1.97; 95% CI= 1.20–3.23), although this study was underpowered for the CRC phenotype (corrected P=0.18; for 25 tested cancer types)^41^. Given these underpowered and contradicting observations, the most recent NCCN guidelines for genetic and familial CRC syndromes (version 2.2017; released on August 9, 2017) concluded that the evidence supporting *ATM* as a CRC-risk gene is deficient and that the risk of CRC in *ATM* mutation carriers is largely unknown^12^. This is the first association study, to our knowledge, that confirmed and independently validated *ATM* as a moderately-penetrant CRC susceptibility gene, explaining the increased risk of colorectal cancer in around 0.74% of all unselected CRC cases. Furthermore, complete loss of *ATM* as a result of acquired deleterious somatic events suggesting a critical role of *ATM* in the CRC tumorigenesis in individuals with inherited *ATM* haploinsufficiency.

In addition to *ATM*, our analysis showed validated evidence supporting germline mutations in *PALB2* as CRC-risk events. *PALB2* plays a critical role in DNA homologous recombination by recruiting *BRCA2* and *RAD51* to DNA breaks to initiate DNA repair. Germline defects in *PALB2* have been associated with breast and pancreatic cancers^25, 42^. Although germline *PALB2* mutations have been observed in several CRC cohorts, it has been so far unclear wither these events contribute to the CRC risk or they merely represent coincidental findings. So far, there has not been any study to evaluate the role of *PALB2* mutations in CRC cases, hence *PALB2* has not been part of the recent NCCN recommendations (version 2.2017) for germline testing in CRC^12^. Our analysis showed evidence for higher-than-expected germline pathogenic *PALB2* mutation rates in around 0.44% of unselected CRC cases, though this effect was not observed in early-onset CRC cohorts. Although tumors of individuals with germline mutations in *PALB2* did not show biallelic inactivation of the gene, our analysis however was not designed to capture potential pathogenic non-coding variants or epigenetic silencing events. Although *ATM* and *PALB2* may only explain a small fraction the CRC heritability in unselected cases, this represents a 20% increase in the diagnostic yield once these two genes are included.

Both *ATM* and *PALB2* are members of homologous recombination (HR) pathway which restores the integrity of double-strand DNA breaks^43^. Inherited HR gene mutations have long been known to increase the risk of several cancers, including breast, ovarian [MIM: 167000], prostate [MIM: 176807] and pancreatic cancers^23, 44, 45^. Here, we showed evidence that germline pathogenic mutations in the HR pathway genes, in aggregate, confer a relative 60-80% increase in the baseline risk of CRC. In addition, biallelic HR gene inactivation, observed in CRCs with various germline HR gene mutations in this study (particularly *ATM* mutation carriers), suggests new venues to explore targeted therapeutic intervention in CRC cases. Breast, ovarian, and prostate cancers from individuals with germline mutations in canonical HR genes have been shown to have substantial response to poly-ADP ribose polymerase (PARP) inhibitors and platinum-based chemotherapy, compared with mutation-negative individuals^46^^-^^48^. As preclinical studies have shown substantial sensitivity of the HR and *ATM*-deficient CRC cell lines to PARPi and with clinical trials to evaluate the efficacy of PARPi in CRC underway (NCT00912743, NCT02305758, NCT01589419, NCT02921256), universal screening of CRC cases for germline HR mutations may provide very informative data that could expand treatment options for these individuals^49^.

The detection of mutations in actionable DRGs has significant ramifications for the probands and their families. First, these mutations significantly increase the person’s risk of developing cancers other than CRC, for several of which effective screening options are available. Furthermore, identifying such mutations in an individual represents a unique opportunity to screen other family members to identify asymptomatic at-risk individuals and implement early surveillance measures. In total, our study estimates that approximately 2.1% (95% CI= 1.1%-3.4%; Binomial Exact) of all CRC cases carry actionable mutations in genes that have not been previously associated with increased CRC risk, which is significantly higher than the combined rate of these mutations in cancer-free controls. In addition, this small but significant subset of CRC cases are, as a result of being carriers of these mutations, at a substantially higher risk of developing several cancers other than CRC. Importantly, these actionable genes are not part of the recommended germline testing for individuals with CRC^12^. Consistent with prior observations in other tumor types, our analysis also demonstrated that positive family history of CRC or other malignancies could not be used as a proxy for the presence of germline DRGs mutations, emphasizing the potential for broader molecular testing strategies to capture these clinically actionable events^50^.

Offering clinical germline molecular testing to cancer cases to evaluate for an inherited cancer-predisposition syndrome relies heavily on several factors such as the individual’s age of presentation and the presence of positive family history of cancer. Intriguingly, our analysis of large CRC cohorts showed that these factors may not reliably predict the likelihood of identifying a germline cancer predisposition mutation in individuals with CRC. First, except for individuals with germline high penetrance CRC risk mutations, our study showed that CRC individuals with low-penetrance CRC risk mutations and those with germline mutations in *ATM* or *PALB2* were not more likely to present at an earlier age compared with presumed sporadic cases. In addition, our study showed that positive family history of CRC was not more commonly reported in CRC individuals who carried high-penetrance CRC risk mutations, low-penetrance CRC risk mutations or DNA repair gene mutations. This is consistent with prior similar observations in the prostate and pediatric cancer spaces^50, 51^. These findings underscore the importance of considering the possibility of carrying an inherited CRC-risk mutation in individuals with late-onset CRC as well as in those without strong family history of CRC. In addition, these observations are also relevant when evaluating the potential utility of implementing early CRC screening measures. However, larger studies are still needed to further delineate the penetrance of these germline mutations.

Our study has several limitations. First, although we performed population stratification, our cases and controls did not come from the same cohort, so enrichment of mutations secondary to non-CRC related factors cannot be completely ruled out. Also, since the raw sequencing data of the control cohort (ExAC) are not publically available, germline variants in cases and controls were not jointly called to limit potential sequencing or pipeline-related variant calling biases. We, however, mitigated this potential source of bias by using the same parameters, tools and platforms that were used to analyze the ExAC cohort. In addition, individual-level clinical information on our control group as well as the validation sets were not available which limited our ability to correct for potential confounders. However, evaluating several independent CRC cohorts makes it unlikely for a confounder to be shared across all cohorts. Finally, larger case-control studies are still necessary to confirm these clinically-relevant findings and inform future updates of clinical germline testing guidelines in CRC cases.

Broadly, our study of large CRC cohorts showed enrichment of disruptive germline pathogenic mutations in the homologous recombination pathway, suggesting its important role in CRC susceptibility and management. In addition, we presented evidence to support *ATM* and *PALB2* as new CRC susceptibility genes, explaining the missing CRC heritability in 1.2% of unselected CRC cases. We also illustrated that a relatively large proportion of all CRC cases have germline pathogenic mutations in HR genes, which may greatly impact their clinical care and inform molecularly driven treatment strategies for individuals with mutations in these genes. Finally, since these genes are not routinely tested clinically, these results could inform revisions to CRC testing guidelines.

## Acknowledgements

We thank all individuals who participated in this study. We also thank Dr. Michele Hacker for her advice on the statistical analysis of this study. We would also like to thank the participants and staff of the NHS and HPFS for their valuable contributions as well as the following state cancer registries for their help: AL, AZ, AR, CA, CO, CT, DE, FL, GA, ID, IL, IN, IA, KY, LA, ME, MD, MA, MI, NE, NH, NJ, NY, NC, ND, OH, OK, OR, PA, RI, SC, TN, TX, VA, WA, WY. Drs. Van Allen, Ogino, Garraway and Fuchs had full access to all the data in the study and take responsibility for the integrity of the data and the accuracy of the data analysis.

This work was conducted with support from Harvard Catalyst, the Harvard Clinical and Translational Science Center (National Center for Research Resources and the National Center for Advancing Translational Sciences, National Institutes of Health Award UL1 TR001102) and financial contributions from Harvard University and its affiliated academic healthcare centers. The content is solely the responsibility of the authors and does not necessarily represent the official views of Harvard Catalyst, Harvard University and its affiliated academic healthcare centers, or the National Institutes of Health. This work was also supported by K08 CA188615-02 (E.M.V.), Damon Runyon Clinical Investigator Award (E.M.V.), KL2 TR001100 (M.G.), R01CA169141-04 (C.S.F.), R01CA118553-07 (C.S.F.), P01 CA87969; UM1 CA186107; P01 CA55075; UM1 CA167552, R35 CA197735 (S.O) and the Stand Up to Cancer Colorectal Cancer Dream Team Translational Research Grant (Grant Number SU2C-AACR-DT22-17) (M.G., C.S.F.). Stand Up to Cancer is a program of the Entertainment Industry Foundation and the research grant is administered by the American Association for Cancer Research, a scientific partner of SU2C. N.D.M. is a Howard Hughes Medical Institute Medical Research Fellow. The funding organizations were not responsible for design and conduct of the study; collection, management, analysis, and interpretation of the data; preparation, review, or approval of the manuscript; and decision to submit the manuscript for publication.

## Availability of data and materials

All BAM files of the CanSeq study are deposited in dbGap phs001075.v1.p1. All raw sequencing files of the NHS/HPFS study are deposited in dbGap phs000722. The TCGA data is available from the database of Genotypes and Phenotypes (dbGaP), Study Accession: phs000178.v9.p8. Raw sequencing data of the NSCCG were not available for analysis, though downstream variant data can be accessed from the “CanVar browser” (https://canvar.icr.ac.uk/).

## Ethics approval and consent to participate

All individuals in the CanSeq study consented to an institutional review board-approved protocol that allows comprehensive genetic analysis of tumor and germline samples (Dana-Farber Cancer Institute #12-078). The NHS/HPFS study was approved by the Partners (IRB#2012-P000788). This study conforms to the Declaration of Helsinki.

## Web Resources section

Online Mendelian Inheritance in Man (http://www.omim.org). <u>Exome Aggregation Consortium</u> (http://exac.broadinstitute.org/). The Cancer Variation Resource (https://canvar.icr.ac.uk/). ClinVar (https://www.ncbi.nlm.nih.gov/clinvar/).

## Supplementary figures

**Figure S1:**
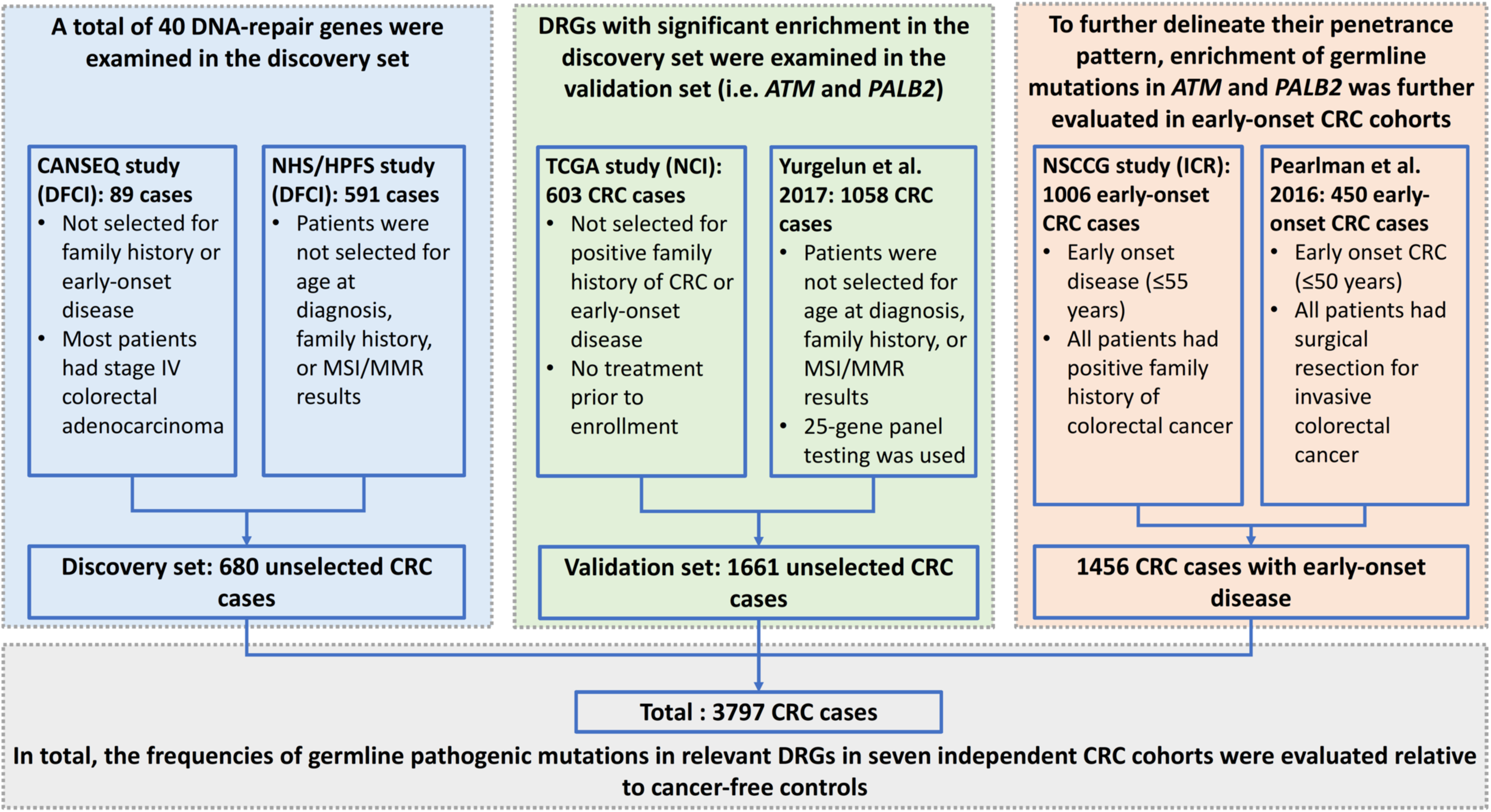
Various cohorts examined in the discovery and validation phases of this study. Two independent cohorts that included 680 CRC individuals were examined in the discovery phase. Of these, a total of 591 CRC cases came from the population-based Nurses’ Health Study (NHS) and the Health Professionals Follow-up Study (HPFS). In addition, 89 CRC cases from the CanSeq study at Dana-Farber Cancer Institute (DFCI) were included in the discovery set. In the validation phase, germline data of 1661 individuals from two independent CRC cohorts were evaluated. Of those, 603 CRC individuals were included in the TCGA project. Individuals in the TCGA cohort were not selected for early-onset disease or positive family history. Germline variants of another 1058 unselected CRC cases who were recently described by Yurgelun et al. were also included in the validation set. Significant findings in the unselected CRC discovery and validation sets were also evaluated in 1456 early-onset CRC cases. In the early-onset CRC set, publically-available germline calls of 1006 early-onset (age<56) familial CRC cases, enrolled in the National Study of Colorectal Cancer Genetics (NSCCG), were examined. Raw sequencing data of the NSCCG were not available for analysis, though downstream variant data was accessed from the “CanVar browser” (https://canvar.icr.ac.uk/; accessed on December 15, 2016). The early-onset CRC set also included 450 CRC individuals who were diagnosed with CRC before the age of 50. The germline variants in these cases were recently described by Pearlman et al, 2017. Raw germline sequencing data of these cohorts were not available for examination. Only germline variants that have been reported in these studies were evaluated. (NHS: Nurses’ Health Study; HPFS: Health Professional Follow Study; TCGA: The Cancer Genome Atlas; NSCCG: National Study of Colorectal Cancer Genetics; ICR: Institute of Cancer Research)

**Figure S2:**
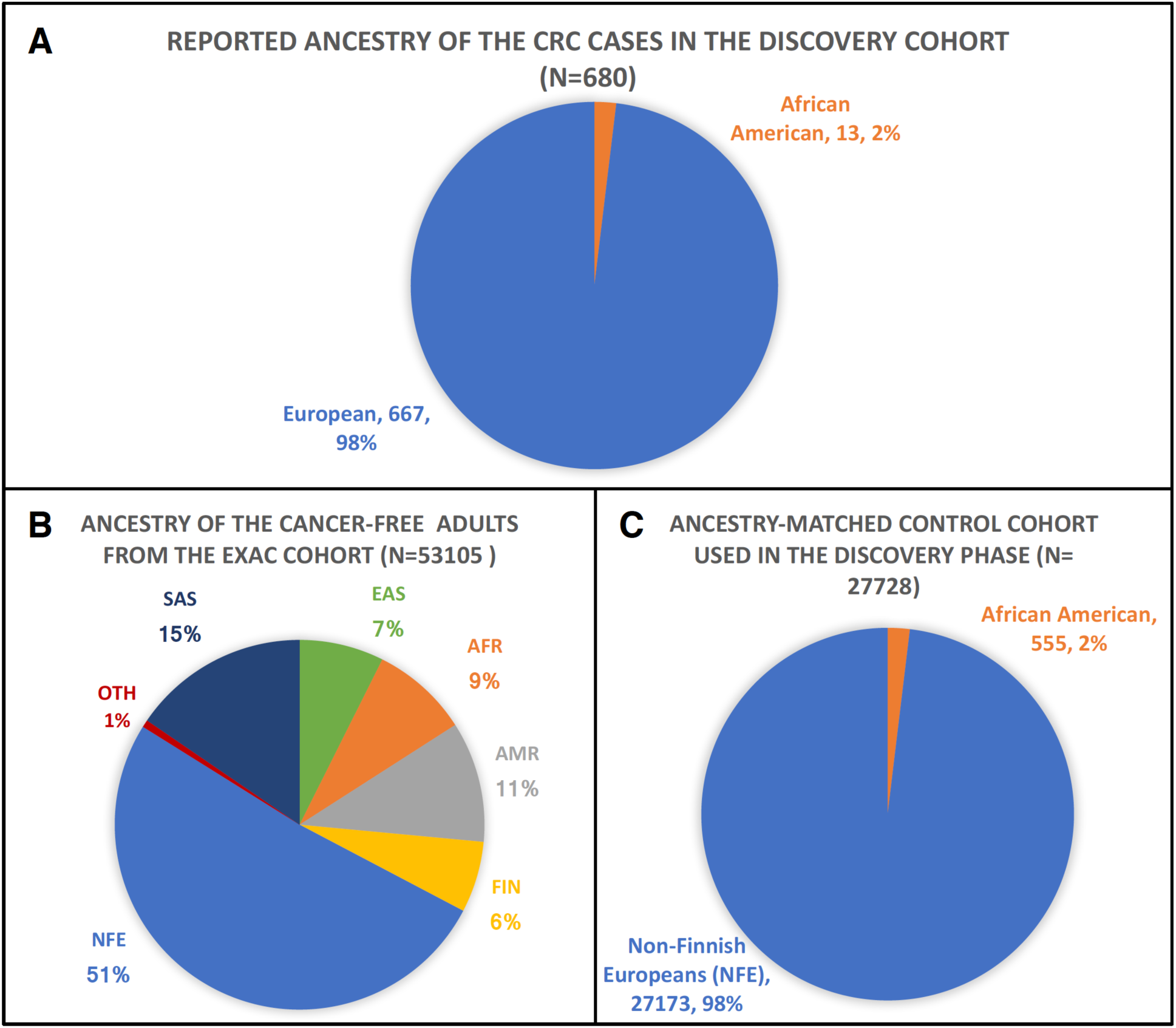
Proportions of cases and controls examined in the discovery phase of this study. A; most of the CRC cases in the discovery set of this study identified their ancestry as European. B&C Rates of germline pathogenic mutations in the examined DRGs were calculated for each of the continental populations reported in the Exome Aggregation Consortium (ExAC) database (African & African American (n=4533), American (n=5608), East Asian (n=3933), Finnish (n=3307), Non-Finnish European (n=27173), South Asian (n=8204)). Based on the proportion of self-reported ancestry representation in our discovery cohort (98% European and 2% African American), ancestry-adjusted frequencies for disruptive mutations in the genes of interest were calculated as follows: Ancestry-adjusted frequency= (0.98 X gene-based frequency of germline pathogenic mutations in NFE) + (0.02 X gene-based frequency of germline pathogenic mutations in AFR). In addition to using ancestry-adjusted rates of mutations as reference values to calculate the significance of enrichment (using Binomial Exact test), we calculated the effect size of enrichment by constructing an ethnicity-matched control cohort (referred to as ExAC_Adj in this study) that constitutes of 27728 individuals (98%; 27173 Non-Finnish Europeans (NFE), and 2%; 555 African Americans (AFR)). Expected number of germline pathogenic mutations in the ancestry-adjusted control cohort in each gene was calculated using the ancestry-adjusted frequency. (AFR: African & African American, AMR: American, EAS: East Asian, FIN: Finnish, NFE: Non-Finnish European, SAS: South Asian, OTH: Other).

**Figure S3:**
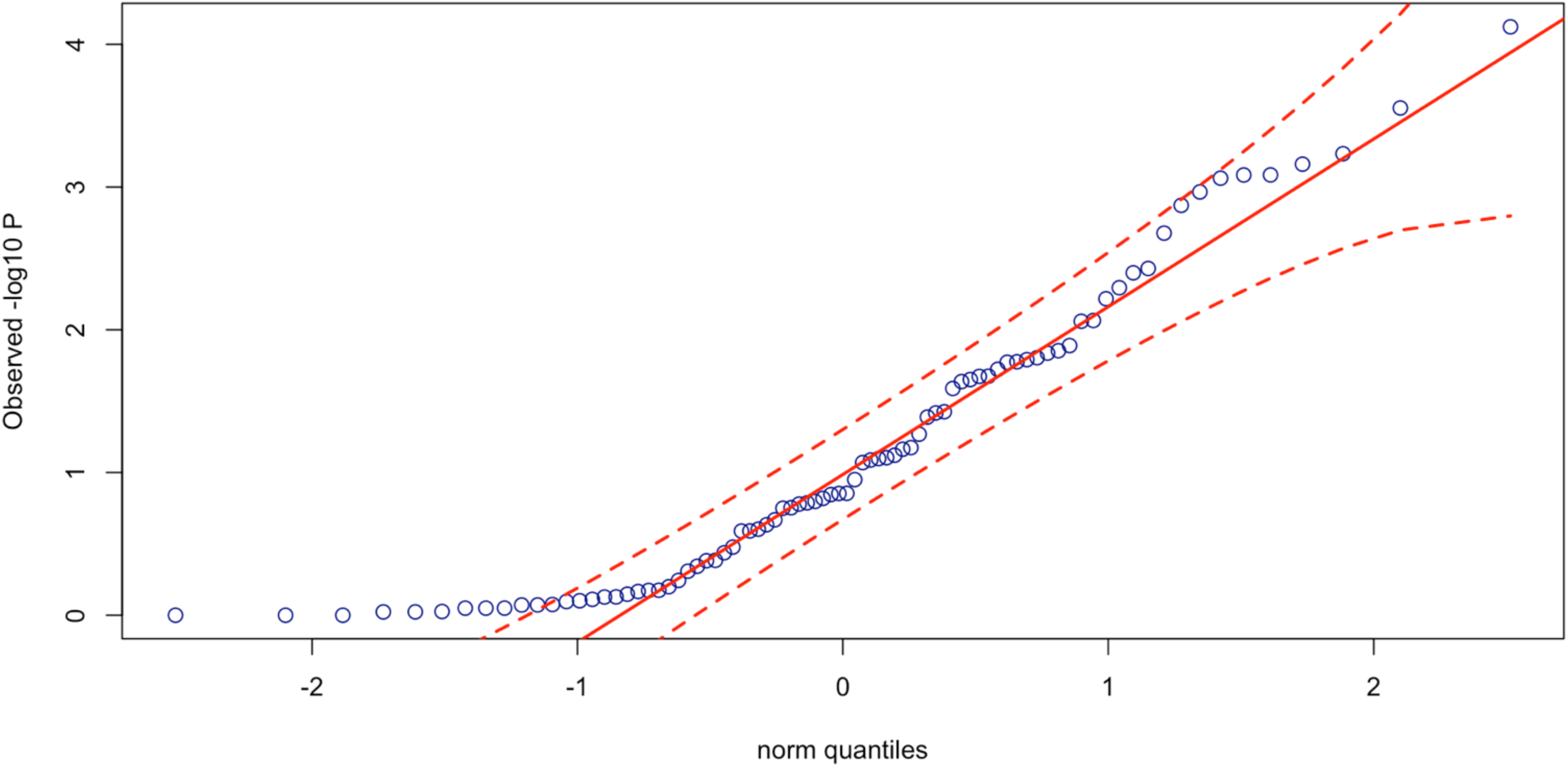
Quantile-quantile plot of the P value of common SNPs in the examined DRGs in the discovery CRC cases compared with the control group (ExAC). No significant deviation from the expected distribution was seen.

**Figure S4:**
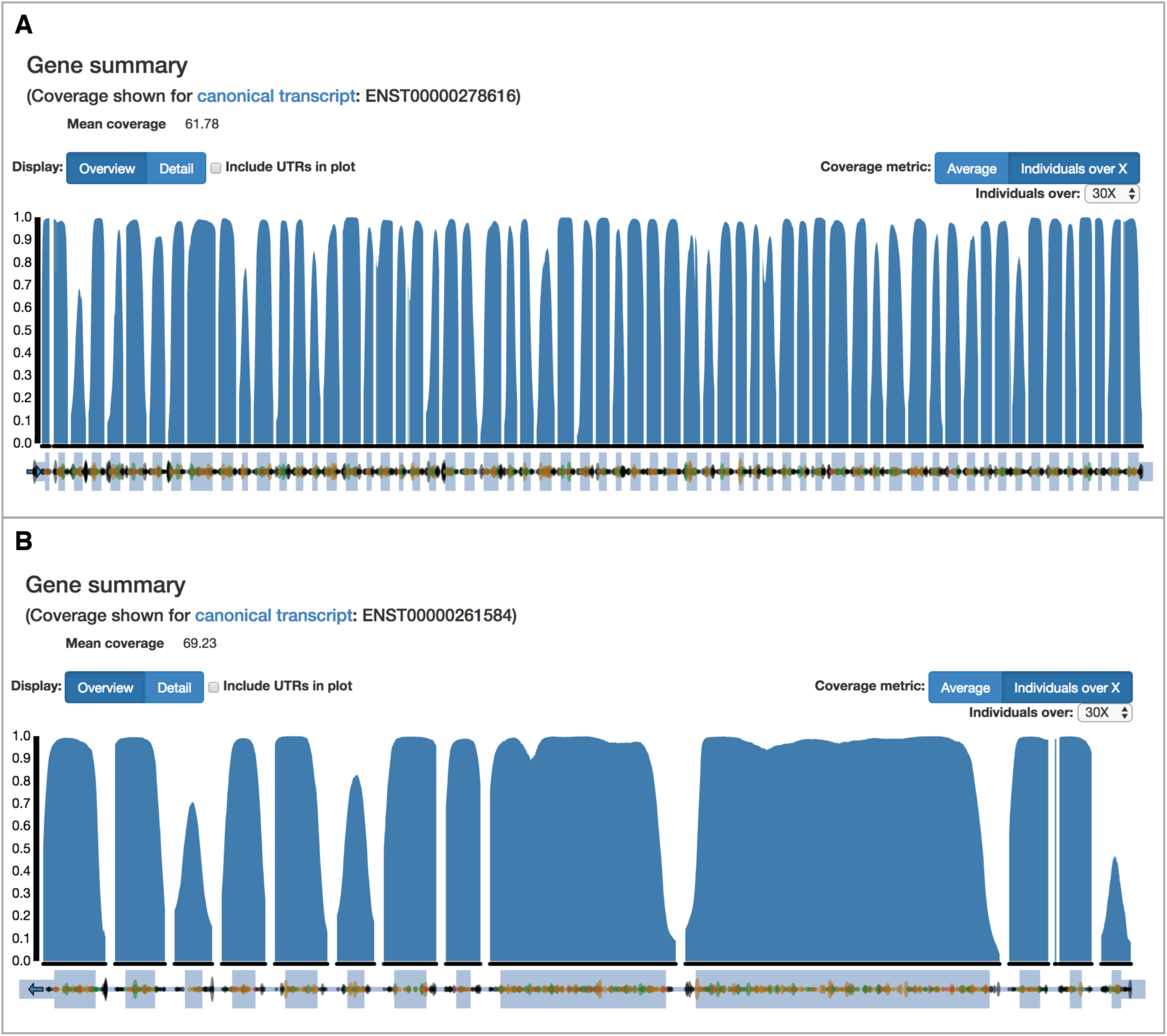
Sequencing coverage of (A) *ATM* and (B) *PALB2* genes in the ExAC cohort, showing the proportion of individuals who had at least 30X coverage for the coding exons.

**Figure S5:**
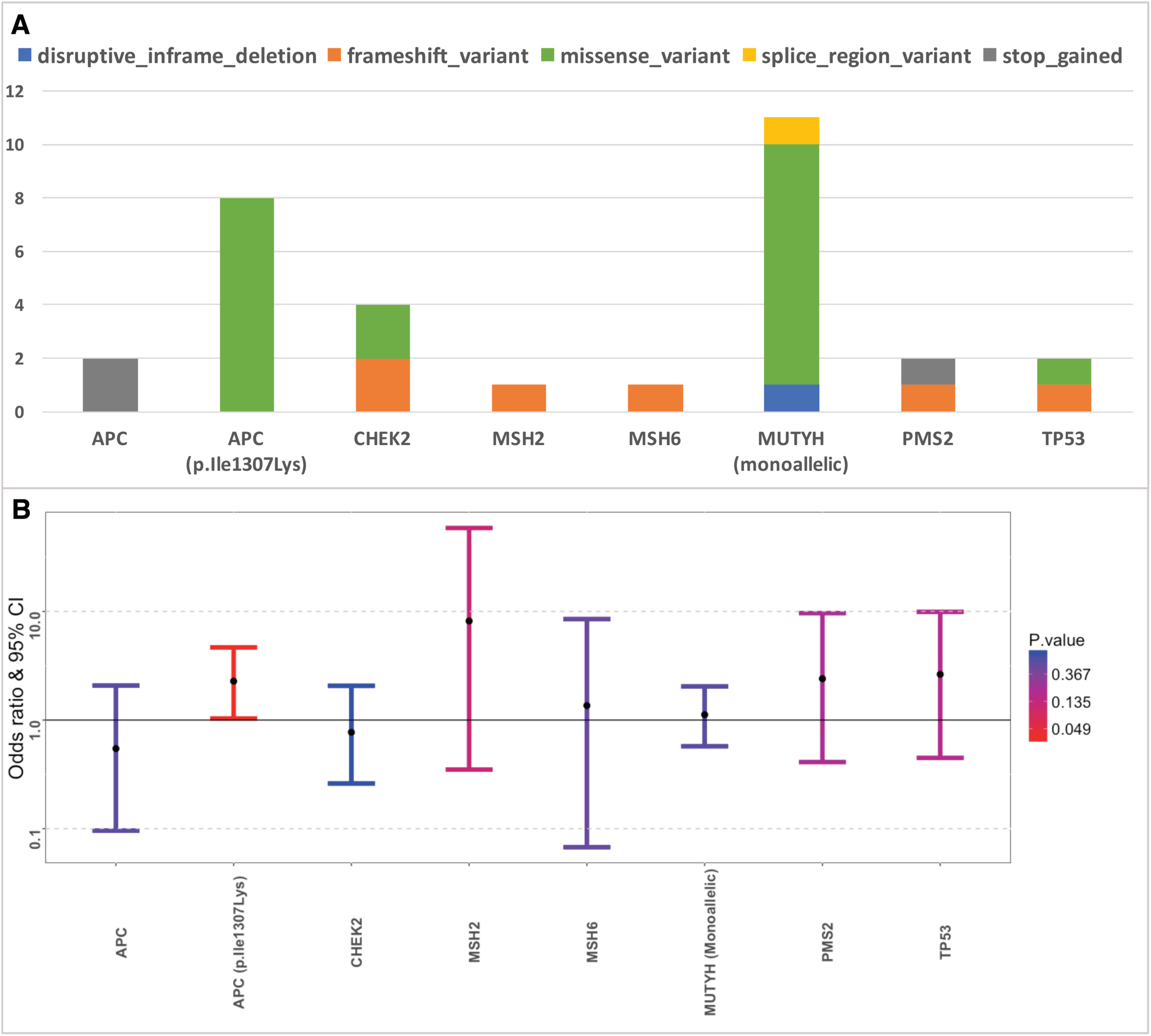
Pathogenic germline mutations in the CRC risk genes in the discovery cohort (n=680). A; Number and impact of detected germline mutations in the examined CRC risk genes. B; Enrichment of germline mutations in the CRC risk genes in the discovery cohort (n=680).

**Figure S6:**
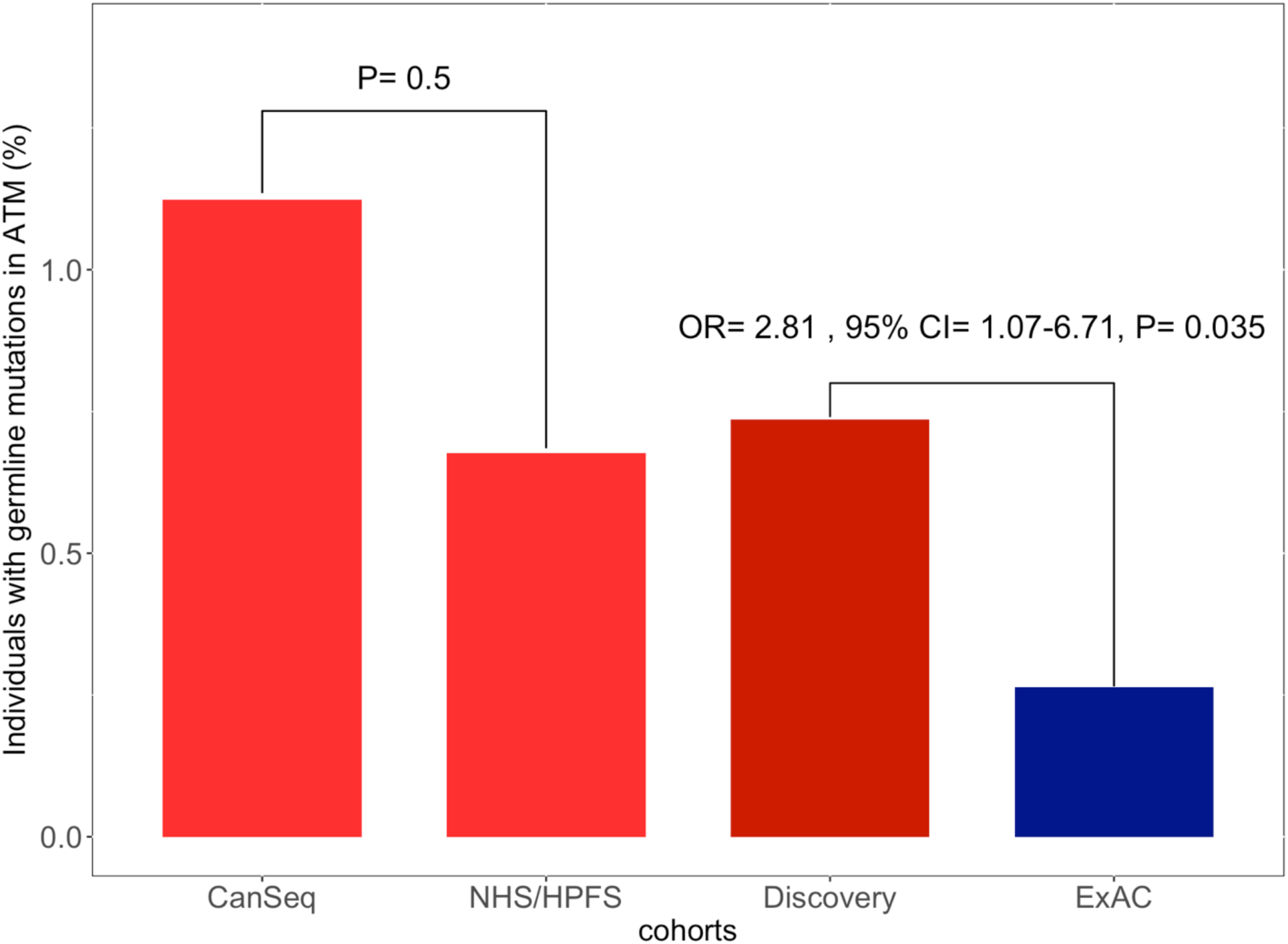
Enrichment of germline pathogenic mutations in *ATM* in each cohort of the discovery set. Our analysis showed that both NHS/HPFS and Canseq cohorts were enriched for *ATM* mutations. There was no statistically significant difference in the frequency of these disruptive events in the Canseq cohort compared with NHS/HPFS (P = 0.5). (NHS: Nurses’ Health Study; HPFS: Health Professional Follow up Study; CanSeq: Cancer Sequencing study)

**Figure S7:**
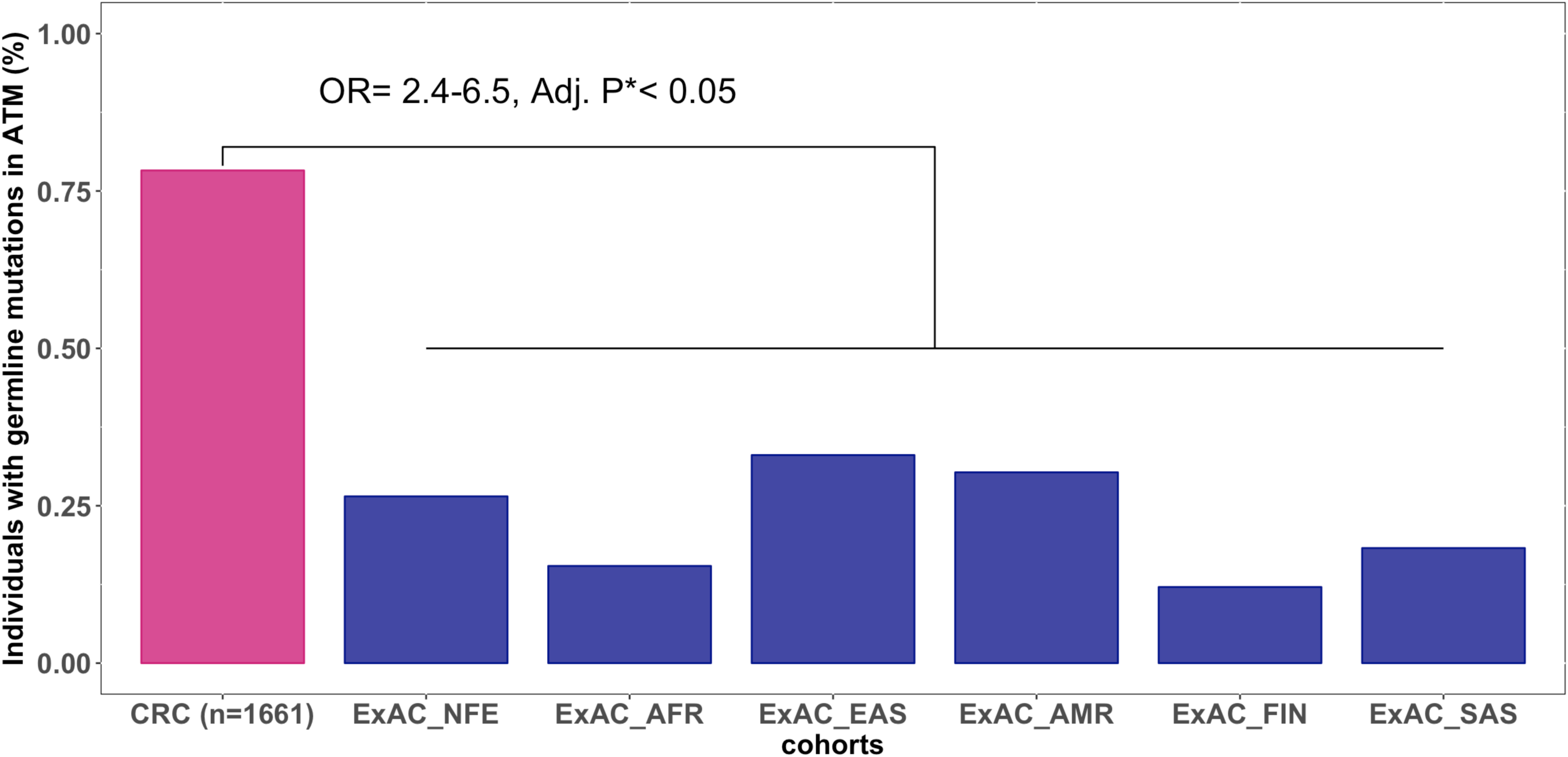
Enrichment of germline *ATM* mutations in the validation set (n= 1661) compared with the various major populations in the ExAC cohort (n=53105; TCGA data excluded; AFR: African & African American, AMR: American, EAS: East Asian, FIN: Finnish, NFE: Non-Finnish European, SAS: South Asian). * P value was adjusted for 6 independent tests using Bonferroni correction

**Figure S8:**
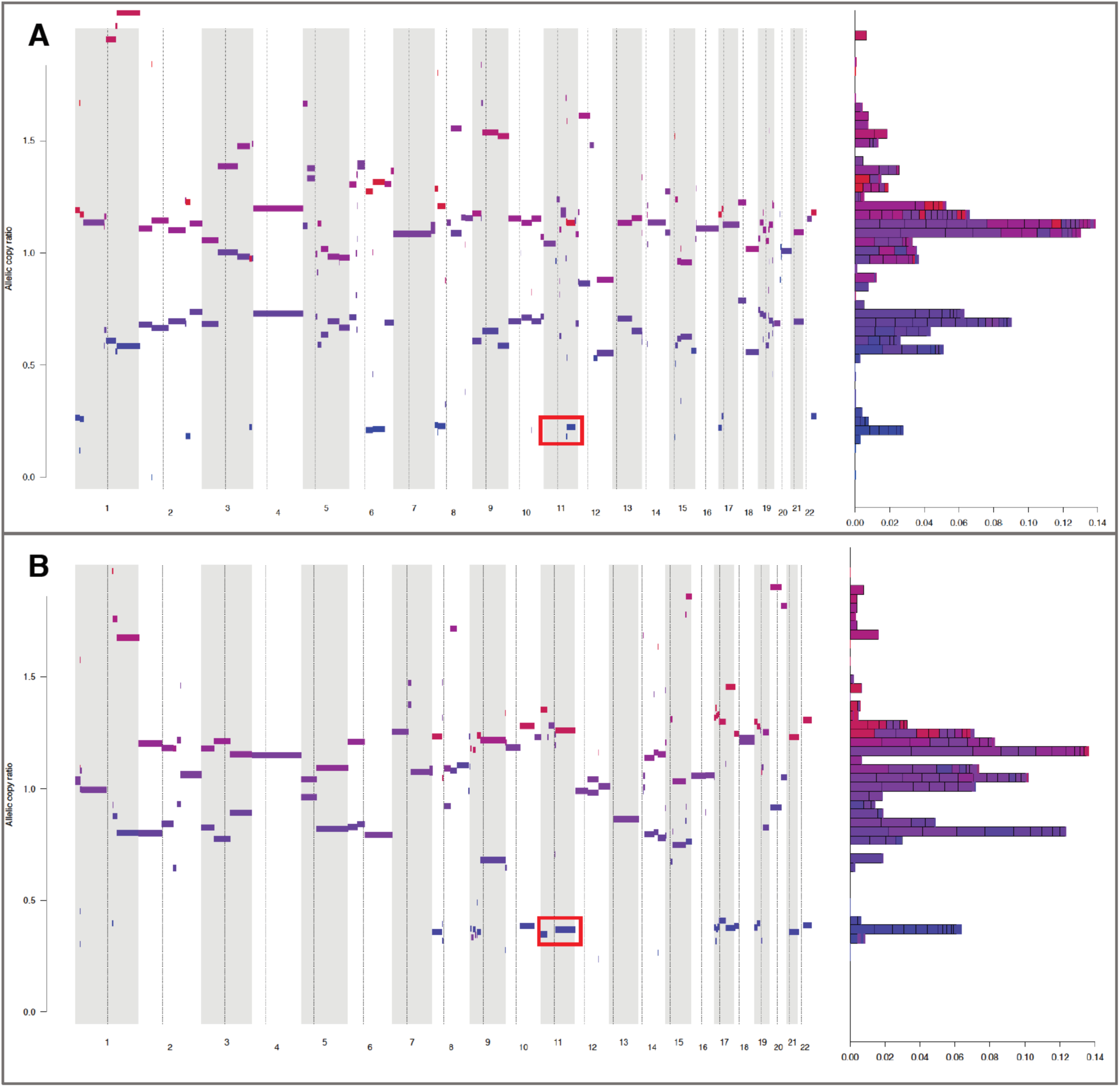
Evaluation of the tumors of cases with germline *ATM* mutations showed LOH of the *ATM* wild-type allele. Two individuals (top: 1221; bottom: 1755) had large deletions involving the cytogenetic region,11q22, which encompasses the *ATM* gene (highlighted).

**Figure S9:**
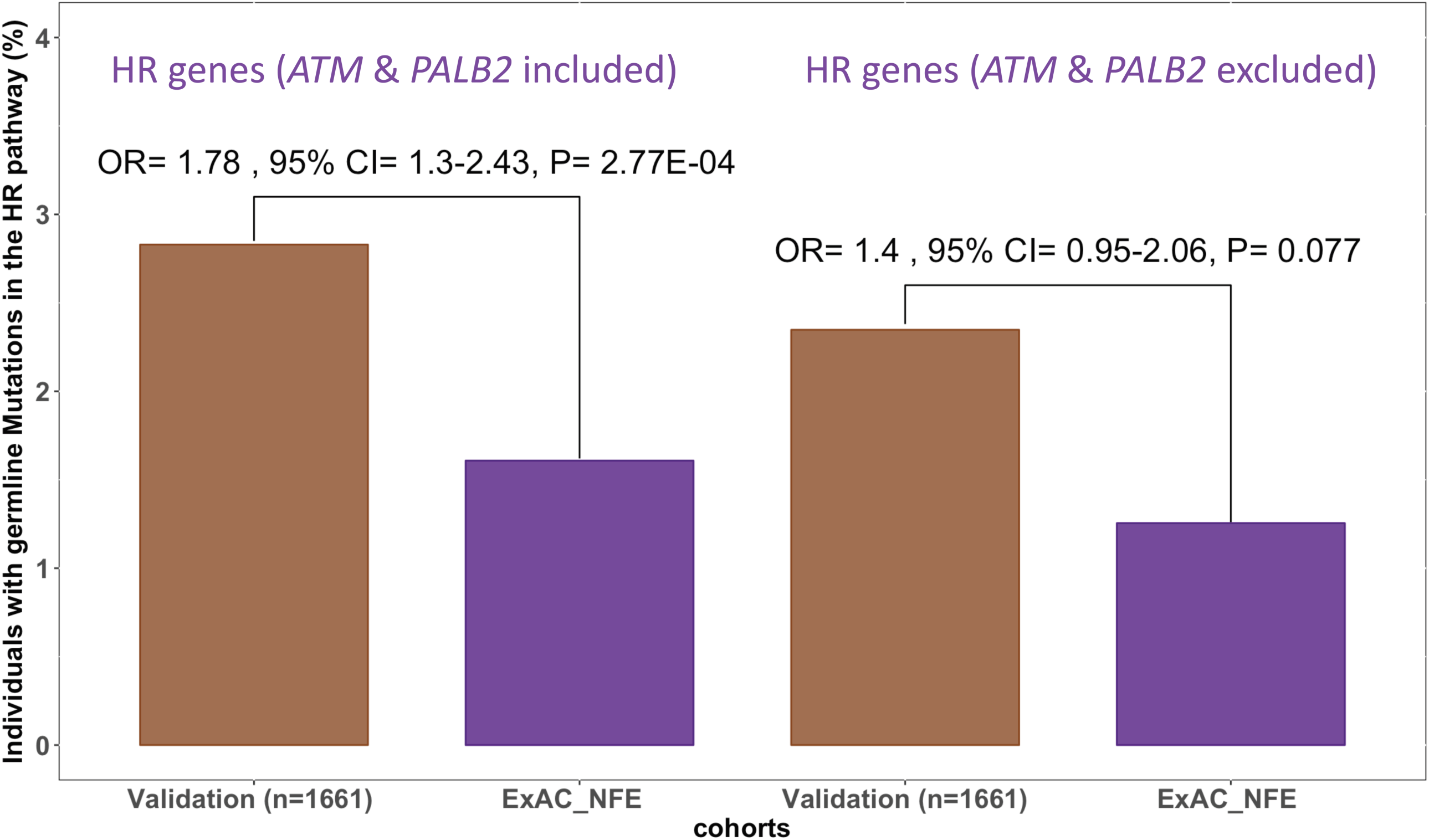
Enrichment of germline pathogenic mutations in the homologous recombination pathway in the CRC validation set. (ExAC: Exome Aggregation Consortium; NFE: Non-Finnish European)

**Figure S10:**
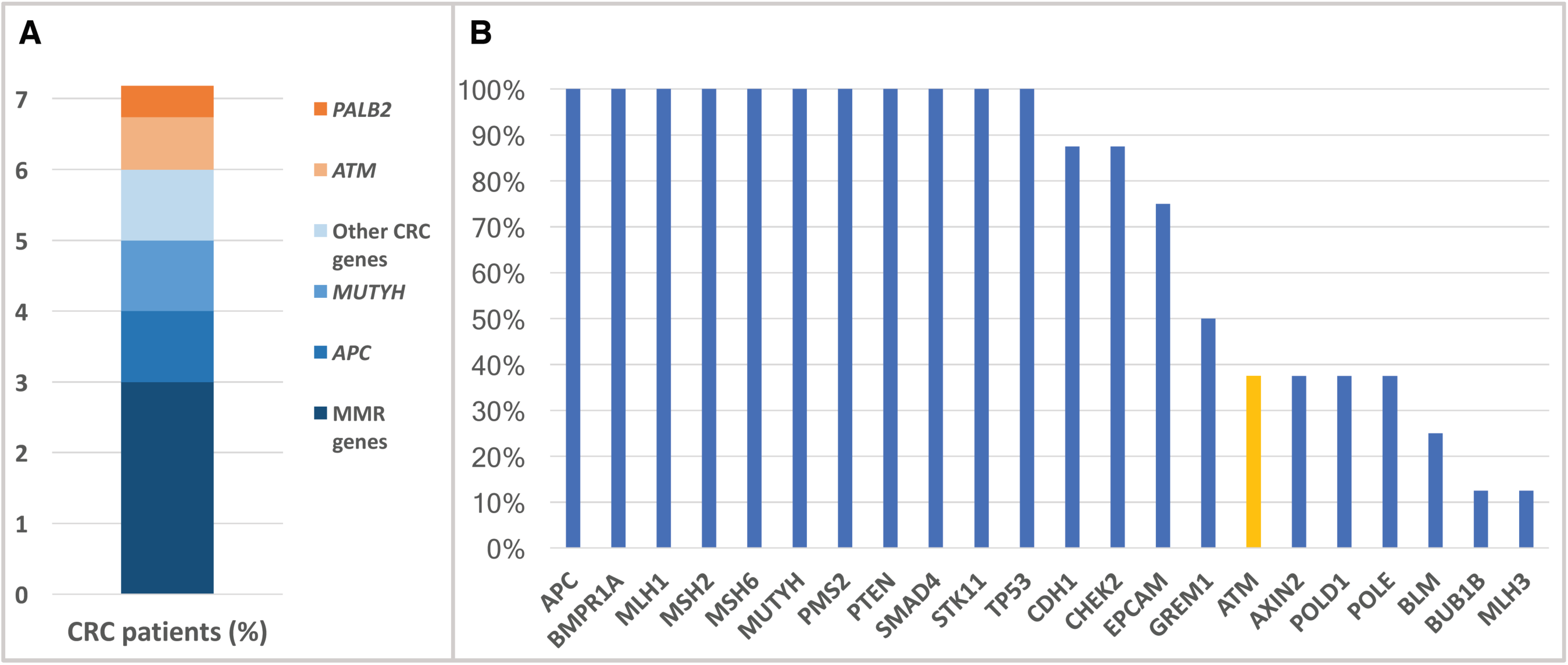
Diagnostic yield of germline testing in unselected CRC cases. A; Although *ATM* and *PALB2* may only explain the CRC heritability in ~1.2% of unselected CRC cases, this represents a potential 20% increase in the current diagnostic yield. B; Genes typically included in the CRC-specific germline testing panels offered by 8 of the largest commercial laboratories in the US (as of August 2017). As shown, *ATM* is only occasionally included in these panels whereas *PALB2* and other highly actionable DRGs are not captured by these clinical tests.

**Figure S11:**
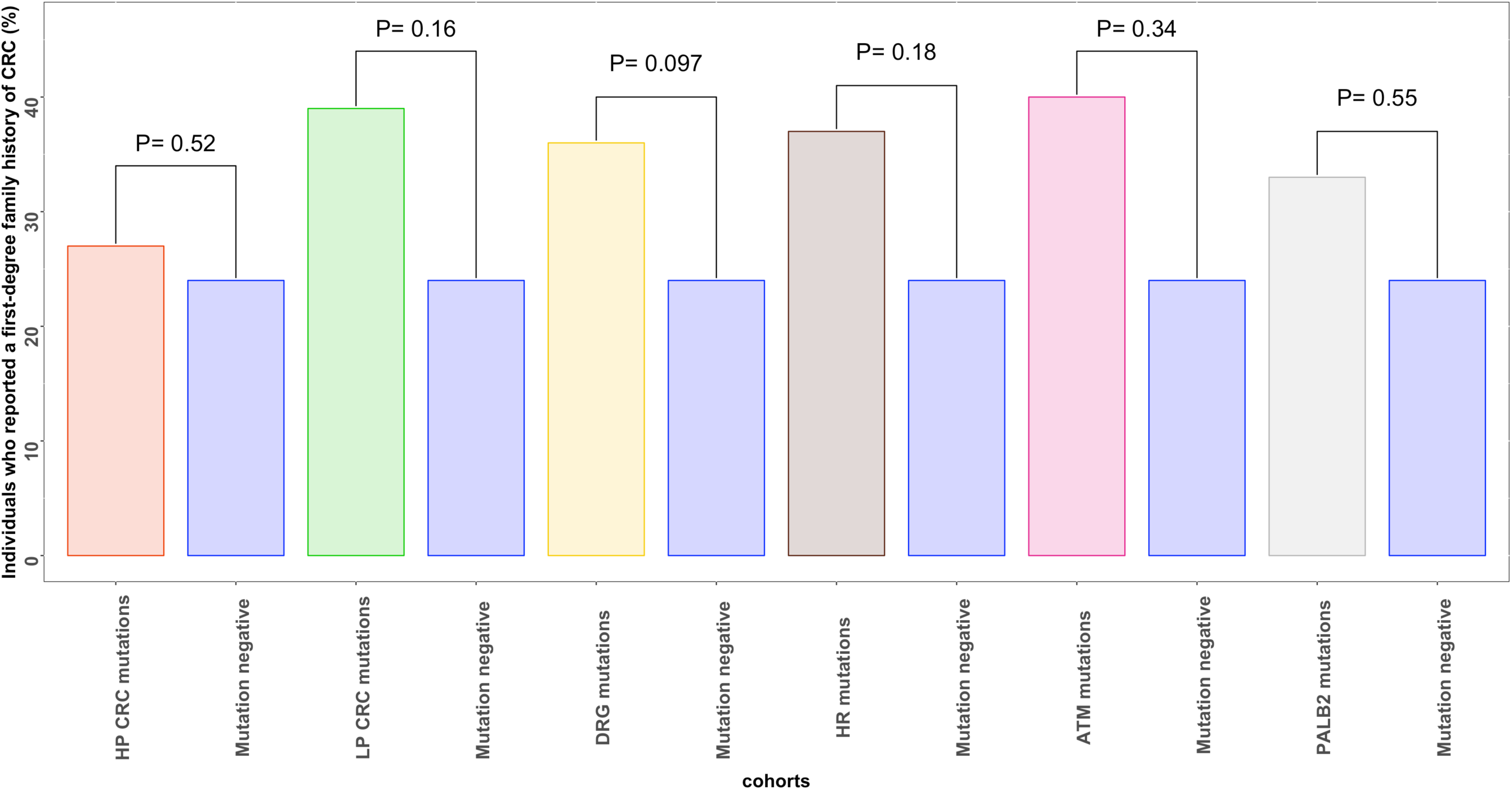
proportions of CRC Individuals who reported positive family history of CRC in one or more first-degree relatives. Individuals with germline pathogenic mutations in the CRC risk genes, DRGs, HR, *ATM* or *PALB2* were not more likely to have a positive family of CRC. Genes contained in each set are listed in Tables 1, S2, and S3.

## Supplementary tables

**Table S1:**
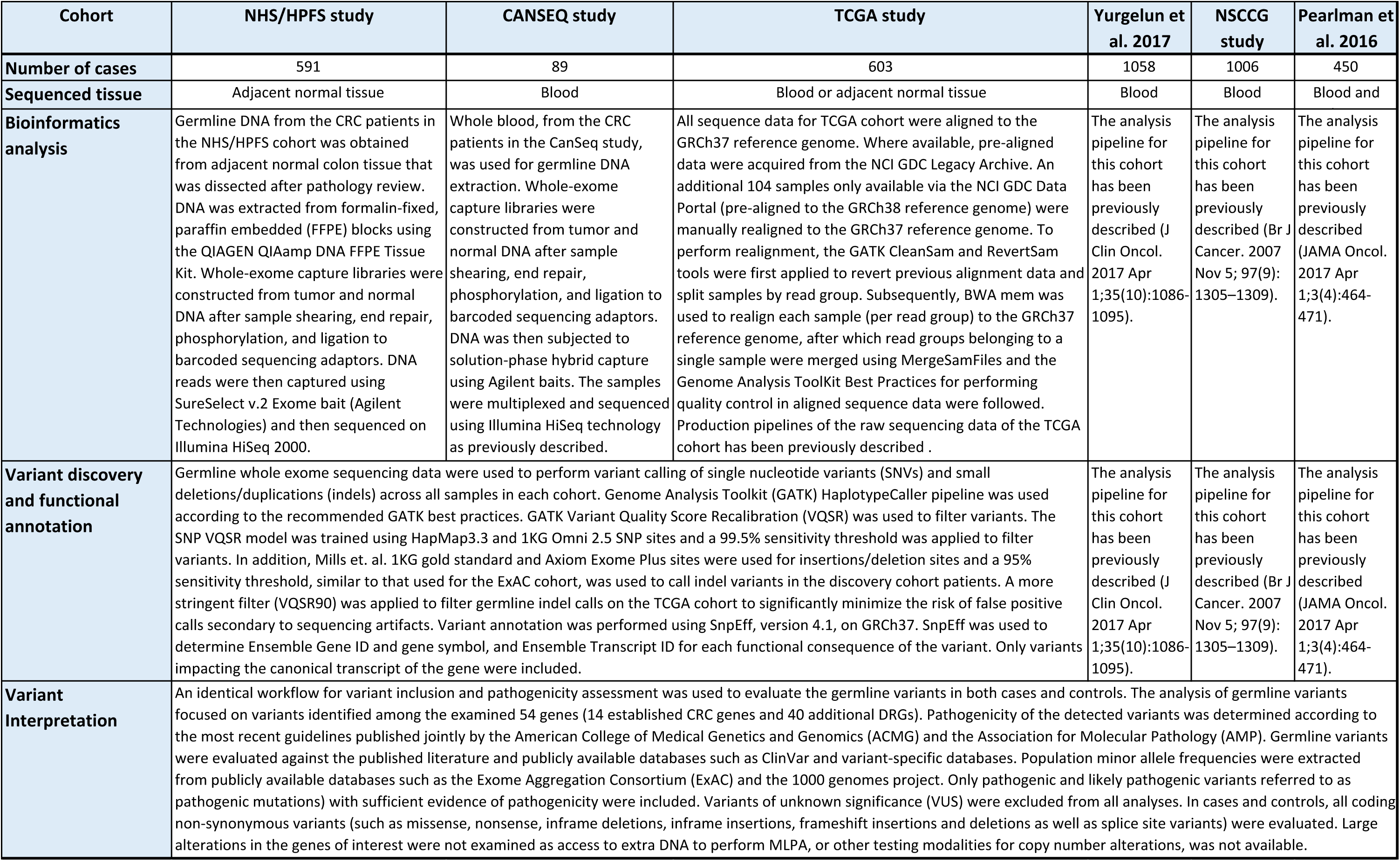
The germline analysis workflows for the examined CRC cohorts in our study.

**Table S2:** DNA repair genes that were evaluated in this study.

| Gene | HGNC Approved Name | Cytogenetic region | Cancer Predisposition Syndrome |
| --- | --- | --- | --- |
| **ATM** | ATM serine/threonine kinase | 11q22-q23 | Ataxia Telangiectasia |
| **ATR** | ATR serine/threonine kinase | 3q23 | Other Cancer Predisposition |
| **BAP1** | BRCA1 associated protein-1 (ubiquitin carboxy-terminal hydrolase) | 3p21.1 | Melanocytic Tumor syndrome, Familial Uveal Melanoma |
| **BARD1** | BRCA1 associated RING domain 1 | 2q35 | Other Cancer Predisposition |
| **BLM** | Bloom syndrome, RecQ helicase-like | 15q26.1 | Bloom Syndrome |
| **BRCA1** | breast cancer 1, early onset | 17q21.31 | Hereditary Breast and Ovarian Cancer |
| **BRCA2** | breast cancer 2, early onset | 13q12-q13 | Hereditary Breast and Ovarian Cancer |
| **BRIP1** | BRCA1 interacting protein C-terminal helicase 1 | 17q22.2 | Other Cancer Predisposition |
| **DDB2** | damage-specific DNA binding protein 2, 48kDa | 11p12-p11 | Xeroderma Pigmentosa |
| **ERCC2** | excision repair cross-complementation group 2 | 19q13.3 | Xeroderma Pigmentosa |
| **ERCC3** | excision repair cross-complementation group 3 | 2q21 | Xeroderma Pigmentosa |
| **ERCC4** | excision repair cross-complementation group 4 | 16p13.3 | Xeroderma Pigmentosa |
| **ERCC5** | excision repair cross-complementation group 5 | 13q22-q34 | Xeroderma Pigmentosa |
| **FANCA** | Fanconi anemia, complementation group A | 16q24.3 | Fanconi Anemia |
| **FANCB** | Fanconi anemia, complementation group B | Xp22.2 | Fanconi Anemia |
| **FANCC** | Fanconi anemia, complementation group C | 9q22.3 | Fanconi Anemia |
| **FANCD2** | Fanconi anemia, complementation group D2 | 3p25.3 | Fanconi Anemia |
| **FANCE** | Fanconi anemia, complementation group E | 6p22-p21 | Fanconi Anemia |
| **FANCF** | Fanconi anemia, complementation group F | 11p15 | Fanconi Anemia |
| **FANCG** | Fanconi anemia, complementation group G | 9p13 | Fanconi Anemia |
| **FANCI** | Fanconi anemia, complementation group I | 15q26.1 | Fanconi Anemia |
| **FANCL** | Fanconi anemia, complementation group L | 2p16.1 | Fanconi Anemia |
| **FANCM** | Fanconi anemia, complementation group M | 14q21.3 | Fanconi Anemia |
| **GEN1** | Holliday junction 5' flap endonuclease | 2p24.2 | Other Cancer Predisposition |
| **MRE11** | MRE11 homolog, double strand break repair nuclease | 11q21 | Ataxia-Telangiectasia-Like Disorder |
| **NBN** | nibrin | 8q21-q24 | Nijmegen Breakage Syndrome |
| **NTHL1** | nth like DNA glycosylase 1 | 16p13.3 | Familial adenomatous polyposis 3 |
| **PALB2** | partner and localizer of BRCA2 | 16p12.1 | Fanconi Anemia |
| **PCNA** | proliferating cell nuclear antigen | 20p12.3 | Ataxia-telangiectasia-like disorder |
| **RAD51** | RAD51 recombinase | 15q15.1 | Breast cancer |
| **RAD51C** | RAD51 paralog C | 17q25.1 | Ovarian cancer |
| **RAD51D** | RAD51 paralog D | 17q11 | Ovarian cancer |
| **RAD54L** | RAD54 like | 1p34.1 | Breast cancer |
| **RECQL4** | RecQ protein-like 4 | 8q24.3 | Rothmund Thomson Syndrome |
| **SLX4** | SLX4 structure-specific endonuclease subunit | 16p13.3 | Fanconi anemia |
| **UBE2T** | ubiquitin conjugating enzyme E2 T | 1q32.1 | Fanconi anemia |
| **WRN** | Werner syndrome, RecQ helicase-like | 8p12 | Werner Syndrome |
| **XPA** | xeroderma pigmentosum, complementation group A | 9q22.3 | Xeroderma Pigmentosa |
| **XPC** | xeroderma pigmentosum, complementation group C | 3p25.1 | Xeroderma Pigmentosa |
| **XRCC3** | X-ray repair cross complementing 3 | 14q32.3 | Breast cancer |

**Table S3:** Established CRC risk genes that were evaluated in this study.

| Gene | HGNC Approved Name | Cytogenetic region | Cancer Predisposition Syndrome |
| --- | --- | --- | --- |
| **APC** | adenomatous polyposis coli | 5q21-q22 | Familial Adenomatous Polyposis |
| **BMPR1A** | bone morphogenetic protein receptor, type IA | 10q22.3 | Hereditary Mixed Polyposis Syndrome |
| **CHEK2** | checkpoint kinase 2 | 22q12.1 | Hereditary Breast |
| **MLH1** | mutL homolog 1 | 3p22.3 | Lynch Syndrome / CMMRD |
| **MSH2** | mutS homolog 2 | 2p21 | Lynch Syndrome / CMMRD |
| **MSH6** | mutS homolog 6 | 2p16 | Lynch Syndrome / CMMRD |
| **MUTYH** | mutY homolog | 1p34.1 | Colorectal cancer |
| **PMS2** | PMS2 postmeiotic segregation increased 2 (S. cerevisiae) | 7p22.1 | Lynch Syndrome / CMMRD |
| **POLD1** | polymerase (DNA directed), delta 1, catalytic subunit | 19q13.3 | Colorectal cancer |
| **POLE** | polymerase (DNA directed), epsilon, catalytic subunit | 12q24.3 | Colorectal cancer |
| **PTEN** | phosphatase and tensin homolog | 10q23 | Cowden syndrome |
| **SMAD4** | SMAD family member 4 | 18q21.1 | Juvenile Polyposis |
| **STK11** | serine/threonine kinase 11 | 19p13.3 | Peutz Jeghers syndrome |
| **TP53** | tumor protein p53 | 17p13.1 | Li Fraumeni Syndrome |

**Table S5:** Depth of sequencing of the examined DRGs in the ExAC cohort.

| Gene | Average depth of coverage (reads) |
| --- | --- |
| *ATM* | 61.78 |
| *ATR* | 58.70 |
| *BAP1* | 56.83 |
| *BARD1* | 58.65 |
| *BLM* | 61.70 |
| *BRCA1* | 66.14 |
| *BRCA2* | 59.19 |
| *BRIP1* | 64.51 |
| *DDB2* | 66.05 |
| *ERCC2* | 50.78 |
| *ERCC3* | 65.46 |
| *ERCC4* | 62.51 |
| *ERCC5* | 58.01 |
| *FANCA* | 52.07 |
| *FANCB* | 60.29 |
| *FANCC* | 48.46 |
| *FANCD2* | 65.88 |
| *FANCE* | 62.03 |
| *FANCF* | 79.42 |
| *FANCG* | 74.03 |
| *FANCI* | 69.40 |
| *FANCL* | 53.48 |
| *FANCM* | 58.54 |
| *GEN1* | 56.12 |
| *MRE11* | 55.41 |
| *NBN* | 60.07 |
| *NTHL1* | 52.13 |
| *PALB2* | 69.23 |
| *PCNA* | 62.64 |
| *RAD51* | 64.25 |
| *RAD51C* | 58.95 |
| *RAD51D* | 50.01 |
| *RAD54L* | 64.53 |
| *RECQL4* | 32.49 |
| *SLX4* | 71.76 |
| *UBE2T* | 67.68 |
| *WRN* | 57.92 |
| *XPA* | 40.62 |
| *XPC* | 45.37 |
| *XRCC3* | 23.71 |
| **Average** | 58.67 |

**Table S6:**
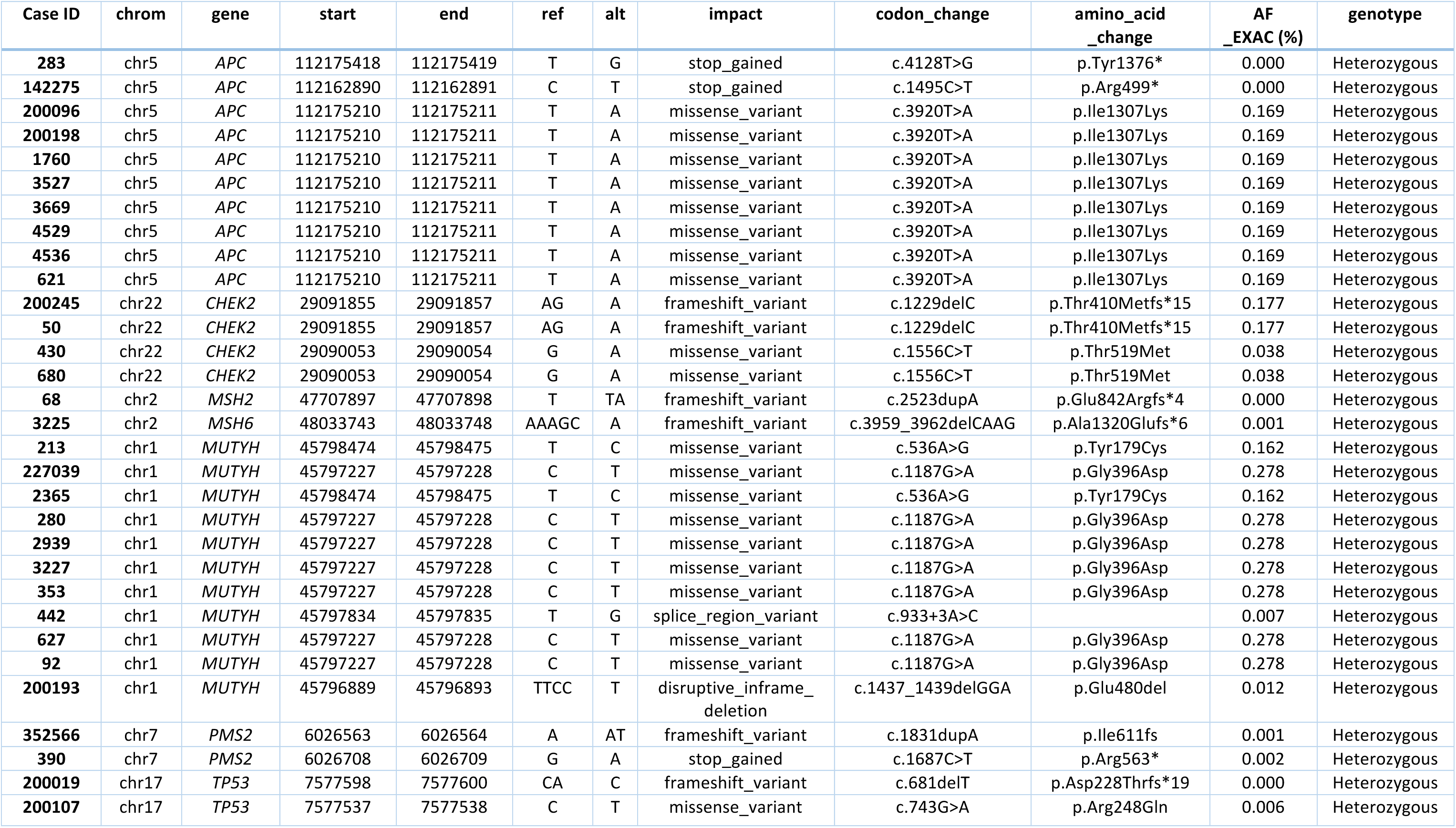
Germline mutations in the well-known CRC risk genes in the CRC discovery set (n=680).

**Table S7:** Germline mutations in the well-known CRC risk genes in the TCGA cohort (n=603).

| Case ID | chrom | gene | start | end | ref | alt | impact | codon_change | amino_acid_change | AF_EXAC (%) | genotype |
| --- | --- | --- | --- | --- | --- | --- | --- | --- | --- | --- | --- |
| TCGA_CRC_18 | chr5 | APC | 112102092 | 112102093 | T | A | stop_gained | c.206T>A | p.Leu69* | 0.001 | Heterozygous |
| TCGA_CRC_01 | chr5 | APC | 112175210 | 112175211 | T | A | missense_variant | c.3920T>A | p.Ile1307Lys | 0.169 | Heterozygous |
| TCGA_CRC_04 | chr5 | APC | 112175210 | 112175211 | T | A | missense_variant | c.3920T>A | p.Ile1307Lys | 0.169 | Heterozygous |
| TCGA_CRC_11 | chr5 | APC | 112175210 | 112175211 | T | A | missense_variant | c.3920T>A | p.Ile1307Lys | 0.169 | Heterozygous |
| TCGA_CRC_16 | chr5 | APC | 112175210 | 112175211 | T | A | missense_variant | c.3920T>A | p.Ile1307Lys | 0.169 | Heterozygous |
| TCGA_CRC_17 | chr5 | APC | 112175210 | 112175211 | T | A | missense_variant | c.3920T>A | p.Ile1307Lys | 0.169 | Heterozygous |
| TCGA_CRC_31 | chr22 | CHEK2 | 29091855 | 29091857 | AG | A | frameshift_variant | c.1229delC | p.Thr410fs | 0.177 | Heterozygous |
| TCGA_CRC_32 | chr22 | CHEK2 | 29091855 | 29091857 | AG | A | frameshift_variant | c.1229delC | p.Thr410fs | 0.177 | Heterozygous |
| TCGA_CRC_33 | chr22 | CHEK2 | 29091855 | 29091857 | AG | A | frameshift_variant | c.1229delC | p.Thr410fs | 0.177 | Heterozygous |
| TCGA_CRC_35 | chr22 | CHEK2 | 29091206 | 29091207 | G | A | missense_variant | c.1412C>T | p.Ser471Phe | 0.030 | Heterozygous |
| TCGA_CRC_12 | chr3 | MLH1 | 37053588 | 37053589 | C | T | stop_gained | c.676C>T | p.Arg226* | 0.001 | Heterozygous |
| TCGA_CRC_03 | chr2 | MSH2 | 47703537 | 47703538 | C | T | stop_gained | c.2038C>T | p.Arg680* | 0.001 | Heterozygous |
| TCGA_CRC_09 | chr2 | MSH2 | 47657023 | 47657024 | T | TC | frameshift_variant | c.1221dupC | p.Tyr408fs | 0.001 | Heterozygous |
| TCGA_CRC_07 | chr2 | MSH6 | 48025862 | 48025864 | AC | A | frameshift_variant | c.742delC | p.Arg248fs | 0.001 | Heterozygous |
| TCGA_CRC_01 | chr1 | MUTYH | 45798465 | 45798466 | C | T | missense_variant | c.545G>A | p.Arg182His | 0.002 | Heterozygous |

**Table S8:**
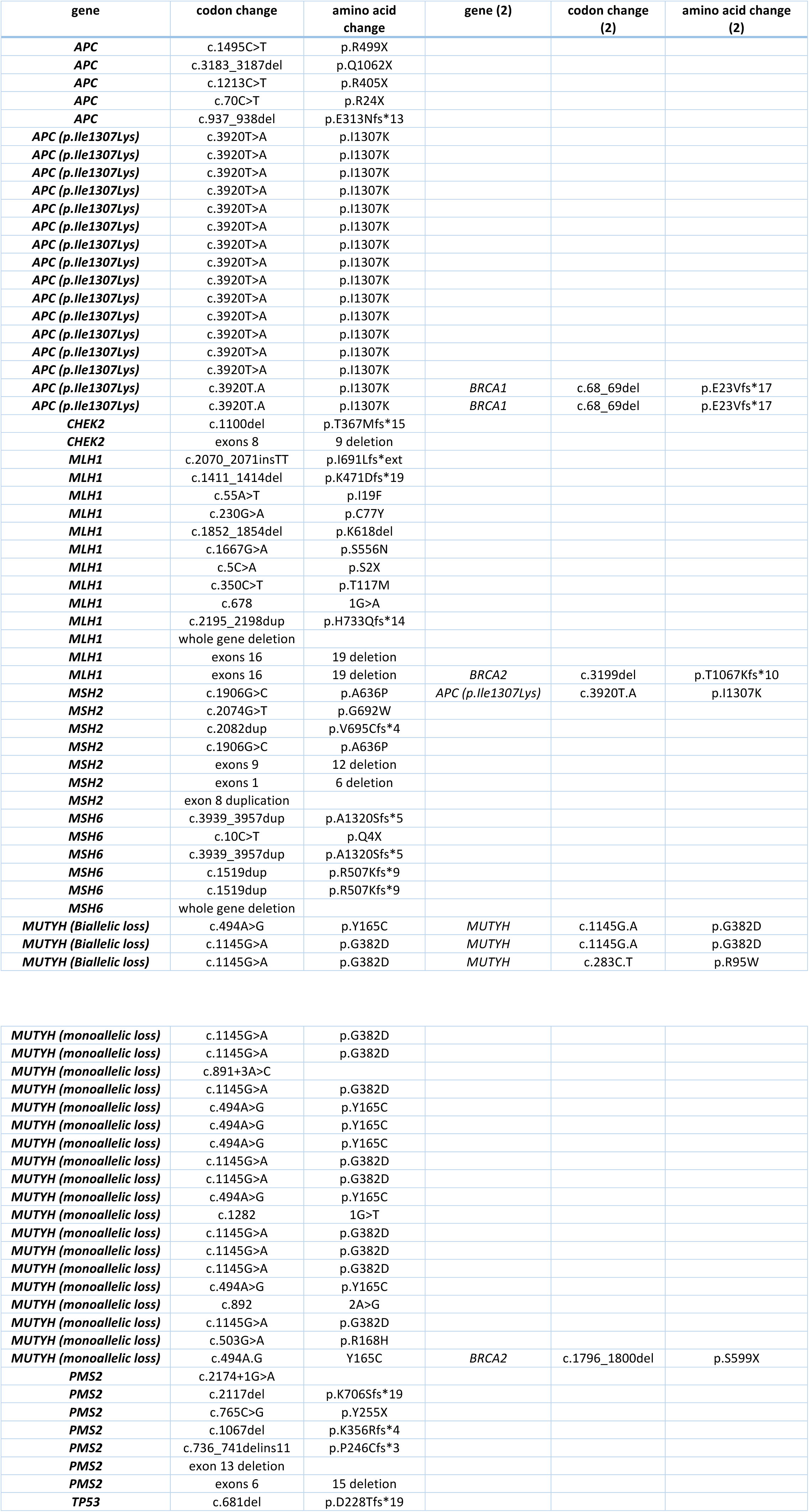
Germline mutations in the well-known CRC risk genes in the Yurgelun et al. 2017 cohort (n=1058).

**Table S9:** Germline mutations in the examined DNA-repair genes in the CRC discovery set (n=680).

| Case ID | chrom | gene | start | end | ref | alt | impact | codon_change | amino_acid_change | AF_EXAC (%) | genotype |
| --- | --- | --- | --- | --- | --- | --- | --- | --- | --- | --- | --- |
| **200205** | chr11 | *ATM* | 108213986 | 108213987 | G | A | stop_gained | c.8307G>A | p.Trp2769* | 0.001 | Heterozygous |
| **1221** | chr11 | *ATM* | 108155006 | 108155008 | AG | A | frameshift_variant | c.3802delG | p.Val1268fs* | 0.003 | Heterozygous |
| **1755** | chr11 | *ATM* | 108141873 | 108141874 | G | T | splice_region_variant | c.2921+1G>T |  | 0.000 | Heterozygous |
| **2760** | chr11 | *ATM* | 108190743 | 108190746 | CAG | C | frameshift_variant | c.6415_6416delGA | p.Glu2139Ilefs*6 | 0.000 | Heterozygous |
| **3645** | chr11 | *ATM* | 108115680 | 108115681 | G | T | stop_gained | c.829G>T | p.Glu277* | 0.000 | Heterozygous |
| **2967** | chr3 | *BAP1* | 52439281 | 52439282 | G | GC | frameshift_variant | c.959dupG | p.Cys320fs | 0.000 | Heterozygous |
| **2775** | chr2 | *BARD1* | 215595201 | 215595204 | CAT | C | frameshift_variant | c.1932_1933delAT | p.Cys645fs | 0.000 | Heterozygous |
| **1743** | chr15 | *BLM* | 91306245 | 91306246 | C | T | stop_gained | c.1933C>T | p.Gln645* | 0.004 | Heterozygous |
| **3046** | chr15 | *BLM* | 91306245 | 91306246 | C | T | stop_gained | c.1933C>T | p.Gln645* | 0.004 | Heterozygous |
| **3181** | chr15 | *BLM* | 91310195 | 91310196 | C | CAAAT | frameshift_variant | c.2250_2251insAAAT | p.Leu751fs | 0.000 | Heterozygous |
| **3111** | chr17 | *BRCA1* | 41209078 | 41209079 | T | TG | frameshift_variant | c.5329dupC | p.Gln1777fs | 0.016 | Heterozygous |
| **200127** | chr13 | *BRCA2* | 32914173 | 32914174 | C | G | stop_gained | c.5682C>G | p.Tyr1894* | 0.000 | Heterozygous |
| **2265** | chr13 | *BRCA2* | 32936731 | 32936732 | G | C | missense_variant | c.7878G>C | p.Trp2626Cys | 0.002 | Heterozygous |
| **2406** | chr13 | *BRCA2* | 32912963 | 32912968 | TGAAA | T | frameshift_variant | c.4478_4481delAAAG | p.Glu1493Valfs*10 | 0.000 | Heterozygous |
| **3444** | chr13 | *BRCA2* | 32890598 | 32890600 | TG | T | frameshift_variant | c.3delG | p.Met1fs | 0.000 | Heterozygous |
| **200054** | chr17 | *BRIP1* | 59761412 | 59761417 | CTTTG | C | frameshift_variant | c.2990_2993delCAAA | p.Thr997Argfs*61 | 0.002 | Heterozygous |
| **204** | chr17 | *BRIP1* | 59885904 | 59885906 | GA | G | frameshift_variant | c.840delT | p.His281Ilefs*8 | 0.000 | Heterozygous |
| **207** | chr19 | *ERCC2* | 45856058 | 45856059 | C | G | missense_variant | c.1847G>C | p.Arg616Pro | 0.013 | Heterozygous |
| **262114** | chr19 | *ERCC2* | 45856058 | 45856059 | C | G | missense_variant | c.1847G>C | p.Arg616Pro | 0.013 | Heterozygous |
| **3558** | chr2 | *ERCC3* | 128050331 | 128050332 | G | A | stop_gained | c.325C>T | p.Arg109* | 0.048 | Heterozygous |
| **3680** | chr16 | *ERCC4* | 14014079 | 14014080 | C | T | stop_gained | c.58C>T | p.Arg20* | 0.000 | Heterozygous |
| **2430** | chr9 | *FANCC* | 97864023 | 97864024 | G | A | stop_gained | c.1642C>T | p.Arg548* | 0.002 | Heterozygous |
| **3439** | chr6 | *FANCE* | 35423605 | 35423607 | GA | G | frameshift_variant | c.334delA | p.Ser112Valfs*14 | 0.000 | Heterozygous |
| **3048** | chr2 | *FANCL* | 58388743 | 58388744 | A | C | stop_gained | c.948T>G | p.Tyr316* | 0.000 | Heterozygous |
| **200226** | chr2 | *GEN1* | 17962406 | 17962411 | TAAAG | T | frameshift_variant | c.1933_1936delAAAG | p.Lys645Cysfs*29 | 0.007 | Heterozygous |
| **251** | chr2 | *GEN1* | 17942842 | 17942845 | AAG | A | frameshift_variant | c.347_348delAG | p.Glu116Valfs*20 | 0.001 | Heterozygous |
| **2760** | chr11 | *MRE11* | 94180441 | 94180442 | G | A | stop_gained | c.1735C>T | p.Arg579* | 0.003 | Heterozygous |
| **1244** | chr11 | *MRE11* | 94200986 | 94200987 | G | A | stop_gained | c.1099C>T | p.Arg367* | 0.005 | Heterozygous |
| **200127** | chr16 | *PALB2* | 23640534 | 23640535 | G | T | stop_gained | c.2576C>A | p.Ser859* | 0.000 | Heterozygous |
| **262114** | chr16 | *PALB2* | 23649451 | 23649452 | T | A | splice_region_variant | c.49-2A>T |  | 0.000 | Heterozygous |
| **587** | chr16 | *PALB2* | 23647355 | 23647358 | ATC | A | frameshift_variant | c.509_510delGA | p.Arg170Ilefs*14 | 0.006 | Heterozygous |
| **101930** | chr6 | *POLH* | 43555063 | 43555064 | G | T | stop_gained | c.328G>T | p.Glu110* | 0.000 | Heterozygous |
| **2946** | chr8 | *RECQL4* | 145741630 | 145741632 | GC | G | frameshift_variant | c.871delG | p.Ala291Leufs*2 | 0.000 | Heterozygous |
| **2957** | chr8 | *RECQL4* | 145738490 | 145738493 | CAT | C | frameshift_variant | c.2492_2493delAT | p.His831Argfs*52 | 0.007 | Heterozygous |
| **2768** | chr16 | *SLX4* | 3641254 | 3641255 | G | C | stop_gained | c.2384C>G | p.Ser795* | 0.000 | Heterozygous |
| **4430** | chr9 | *XPA* | 100459470 | 100459471 | G | GC | frameshift_variant | c.103dupG | p.Ala35Glyfs*27 | 0.000 | Heterozygous |
| **587** | chr14 | *XRCC3* | 104165866 | 104165869 | GAC | G | frameshift_variant | c.606_607delGT | p.Arg204Glyfs*18 | 0.001 | Heterozygous |

**Table S10:** Germline mutations in *ATM*, *PALB2* and other HR genes in the TCGA cohort (n=603).

| Case ID | chrom | gene | start | end | ref | alt | impact | codon_change | aa_change | AF_EXAC (%) | genotype |
| --- | --- | --- | --- | --- | --- | --- | --- | --- | --- | --- | --- |
| TCGA_CRC_02 | chr11 | *ATM* | 108186741 | 108186742 | C | T | stop_gained | c.6100C>T | p.Arg2034* | 0.000 | Heterozygous |
| TCGA_CRC_14 | chr11 | *ATM* | 108205831 | 108205832 | T | C | missense_variant | c.8147T>C | p.Val2716Ala | 0.004 | Heterozygous |
| TCGA_CRC_05 | chr11 | *ATM* | 108224607 | 108224608 | G | A | splice_donor_variant | c.8786+1G>A |  | 0.002 | Heterozygous |
| TCGA_CRC_22 | chr2 | *BARD1* | 215610565 | 215610566 | G | A | stop_gained | c.1690C>T | p.Gln564* | 0.005 | Heterozygous |
| TCGA_CRC_26 | chr15 | *BLM* | 91293264 | 91293267 | ACT | A | frameshift_variant | c.772_773delCT | p.Leu258Glufs*7 | 0.004 | Heterozygous |
| TCGA_CRC_27 | chr15 | *BLM* | 91303903 | 91303904 | C | G | stop_gained | c.1301C>G | p.Ser434* | 0.001 | Heterozygous |
| TCGA_CRC_28 | chr15 | *BLM* | 91304244 | 91304245 | C | T | stop_gained | c.1642C>T | p.Gln548* | 0.018 | Heterozygous |
| TCGA_CRC_29 | chr15 | *BLM* | 91304244 | 91304245 | C | T | stop_gained | c.1642C>T | p.Gln548* | 0.018 | Heterozygous |
| TCGA_CRC_30 | chr15 | *BLM* | 91306245 | 91306246 | C | T | stop_gained | c.1933C>T | p.Gln645* | 0.004 | Heterozygous |
| TCGA_CRC_08 | chr17 | *BRCA1* | 41245089 | 41245091 | TG | T | frameshift_variant | c.2457delC | p.Asp821fs | 0.000 | Heterozygous |
| TCGA_CRC_15 | chr13 | *BRCA2* | 32914436 | 32914438 | GT | G | frameshift_variant | c.5946delT | p.Ser1982fs | 0.026 | Heterozygous |
| TCGA_CRC_34 | chr13 | *BRCA2* | 32936731 | 32936732 | G | C | missense_variant | c.7878G>C | p.Trp2626Cys | 0.002 | Heterozygous |
| TCGA_CRC_10 | chr17 | *BRIP1* | 59793411 | 59793412 | G | A | stop_gained | c.2392C>T | p.Arg798* | 0.015 | Heterozygous |
| TCGA_CRC_23 | chr8 | *NBN* | 90983440 | 90983446 | ATTTGT | A | frameshift_variant | c.657_661delACAAA | p.Lys219fs | 0.019 | Heterozygous |
| TCGA_CRC_24 | chr8 | *NBN* | 90983440 | 90983446 | ATTTGT | A | frameshift_variant | c.657_661delACAAA | p.Lys219fs | 0.019 | Heterozygous |
| TCGA_CRC_25 | chr8 | *NBN* | 90983440 | 90983446 | ATTTGT | A | frameshift_variant | c.657_661delACAAA | p.Lys219fs | 0.019 | Heterozygous |
| TCGA_CRC_20 | chr16 | *PALB2* | 23647107 | 23647108 | T | TA | frameshift_variant | c.758dupT | p.Ser254fs | 0.003 | Heterozygous |
| TCGA_CRC_06 | chr16 | *PALB2* | 23647355 | 23647358 | ATC | A | frameshift_variant | c.509_510delGA | p.Arg170fs | 0.006 | Heterozygous |
| TCGA_CRC_13 | chr16 | *PALB2* | 23647355 | 23647358 | ATC | A | frameshift_variant | c.509_510delGA | p.Arg170fs | 0.006 | Heterozygous |

**Table S11:** Germline mutations in *ATM*, *PALB2* and other HR genes in the Yurgelun et al. 2017 cohort (n=1058).

| gene | codon change | amino acid change | gene (2) | codon change (2) | amino acid change (2) |
| --- | --- | --- | --- | --- | --- |
| **BRCA1** | c.68_69del | p.E23Vfs*17 | *APC (p.Ile1307Lys)* | c.3920T.A | p.I1307K |
| **BRCA1** | c.68_69del | p.E23Vfs*17 | *APC (p.Ile1307Lys)* | c.3920T.A | p.I1307K |
| **BRCA2** | c.3199del | p.T1067Kfs*10 | *MLH1* | exons 16-19 deletion |  |
| **BRCA2** | c.1796_1800del | p.S599X | *MUTYH (monoallelic loss)* | c.494A.G | Y165C |
| **ATM** | c.8934_8935del | p.E2979Afs*9 |  |  |  |
| **ATM** | c.7638_7646del | p.R2547_S2549del |  |  |  |
| **ATM** | c.4632_4635del | p.Y1544X |  |  |  |
| **ATM** | c.3760del | p.V1254Ffs*2 |  |  |  |
| **ATM** | c.802C>T | p.Q268X |  |  |  |
| **ATM** | c.790del | p.Y264Ifs*12 |  |  |  |
| **ATM** | c.5570C>A | p.S1857X |  |  |  |
| **ATM** | c.2413C>T | p.R805X |  |  |  |
| **ATM** | c.2250G>A | p.K750K |  |  |  |
| **ATM** | c.3480_3492dup | p.S1165Gfs*5 |  |  |  |
| **BRCA1** | c.5095C>T | p.R1699W |  |  |  |
| **BRCA2** | c.7602del | p.C2535Vfs*16 |  |  |  |
| **BRCA2** | c.5946del | p.S1982Rfs*22 |  |  |  |
| **BRCA2** | c.4477G>T | p.E1493X |  |  |  |
| **BRCA2** | c.3847_3848del | p.V1283Kfs*2 |  |  |  |
| **BRCA2** | c.8537_8538del | p.E2846Gfs*22 |  |  |  |
| **BRCA2** | c.8537_8538del | p.E2846Gfs*22 |  |  |  |
| **BRIP1** | c.2990_2993del | p.T997Rfs*61 |  |  |  |
| **BRIP1** | c.2379+1G>T |  |  |  |  |
| **BRIP1** | c.1970del | p.G657Vfs*31 |  |  |  |
| **NBN** | c.657_661del | p.K219Nfs*16 |  |  |  |
| **NBN** | c.1142del | p.P381Qfs*23 |  |  |  |
| **PALB2** | c.2711G>A | p.W904X |  |  |  |
| **PALB2** | c.751C>T | p.Q251X |  |  |  |

**Table S12:** Somatic inactivating mutations presumably affecting the wild-type allele of genes where germline mutations were detected.

| Case ID | gene | germline mutation |  |  | Somatic LOH evaluation |  |  |
| --- | --- | --- | --- | --- | --- | --- | --- |
|  |  | impact | codon_change | aa_change | Large Deletion | Point mutation | LOH Call |
| 283 | APC | stop_gained | c.4128T>G | p.Tyr1376* | unknown | unknown | unknown |
| 142275 | APC | stop_gained | c.1495C>T | p.Arg499* | No | No | No |
| 621 | APC (p.Ile1307Lys) | missense_variant | c.3920T>A | p.Ile1307Lys | Yes | Yes (1 frameshift insertion) | Yes |
| 1760 | APC (p.Ile1307Lys) | missense_variant | c.3920T>A | p.Ile1307Lys | No | Yes (1 nonsense mutation) | Yes |
| 3527 | APC (p.Ile1307Lys) | missense_variant | c.3920T>A | p.Ile1307Lys | No | Yes (1 splice site) | Yes |
| 3669 | APC (p.Ile1307Lys) | missense_variant | c.3920T>A | p.Ile1307Lys | No | Yes (2 frameshift insertion) | Yes |
| 4529 | APC (p.Ile1307Lys) | missense_variant | c.3920T>A | p.Ile1307Lys | Yes | Yes (1 nonsense mutation) | Yes |
| 4536 | APC (p.Ile1307Lys) | missense_variant | c.3920T>A | p.Ile1307Lys | No | Yes (1 nonsense mutation) | Yes |
| 200096 | APC (p.Ile1307Lys) | missense_variant | c.3920T>A | p.Ile1307Lys | No | Yes (1 nonsense mutation, 1 frameshift insertion) | Yes |
| 200198 | APC (p.Ile1307Lys) | missense_variant | c.3920T>A | p.Ile1307Lys | No | Yes (1 nonsense mutation, frameshift deletion) | Yes |
| 1221 | ATM | frameshift_variant | c.3802delG | p.Val1268fs | Yes | No | Yes |
| 1755 | ATM | splice_region_variant | c.2921+1G>T |  | Yes | No | Yes |
| 2760 | ATM | frameshift_variant | c.6415_6416delGA | p.Glu2139Ilefs*6 | No | Yes (1 nonsense mutation) | Yes |
| 3645 | ATM | stop_gained | c.829G>T | p.Glu277* | No | Yes (1 nonsense mutation) | Yes |
| 200205 | ATM | stop_gained | c.8307G>A | p.Trp2769* | No | Yes (1 splice mutation) | Yes |
| 2967 | BAP1 | frameshift_variant | c.959dupG | p.Cys320fs | No | No | No |
| 2775 | BARD1 | frameshift_variant | c.1932_1933delAT | p.Cys645fs | No | No | No |
| 1743 | BLM | stop_gained | c.1933C>T | p.Gln645* | No | No | No |
| 3046 | BLM | stop_gained | c.1933C>T | p.Gln645* | No | No | No |
| 3181 | BLM | frameshift_variant | c.2250_2251insAAAT | p.Leu751fs | No | No | No |
| 3111 | BRCA1 | frameshift_variant | c.5329dupC | p.Gln1777fs | unknown | unknown | unknown |
| 2265 | BRCA2 | missense_variant | c.7878G>C | p.Trp2626Cys | No | No | No |
| 2406 | BRCA2 | frameshift_variant | c.4478_4481delAAAG | p.Glu1493Valfs*10 | No | No | No |
| 3444 | BRCA2 | frameshift_variant | c.3delG | p.Met1fs | No | Yes (1 missense mutation) | Yes |
| 200127 | BRCA2 | stop_gained | c.5682C>G | p.Tyr1894* | No | No | No |
| 204 | BRIP1 | frameshift_variant | c.840delT | p.His281Ilefs*8 | No | No | No |
| 200054 | BRIP1 | frameshift_variant | c.2990_2993delCAAA | p.Thr997Argfs*61 | No | No | No |
| 200245 | CHEK2 | frameshift_variant | c.1229delC | p.Thr410Metfs*15 | unknown | unknown | unknown |
| 50 | CHEK2 | frameshift_variant | c.1229delC | p.Thr410Metfs*15 | Yes | No | Yes |
| 430 | CHEK2 | missense_variant | c.1556C>T | p.Thr519Met | No | No | No |
| 680 | CHEK2 | missense_variant | c.1556C>T | p.Thr519Met | No | No | No |
| 207 | ERCC2 | missense_variant | c.1847G>C | p.Arg616Pro | No | Yes (1 frameshift deletion) | Yes |
| 262114 | ERCC2 | missense_variant | c.1847G>C | p.Arg616Pro | Yes | No | Yes |
| 3558 | ERCC3 | stop_gained | c.325C>T | p.Arg109* | No | No | No |
| 3680 | ERCC4 | stop_gained | c.58C>T | p.Arg20* | No | No | No |
| 2430 | FANCC | stop_gained | c.1642C>T | p.Arg548* | No | No | No |
| 3439 | FANCE | frameshift_variant | c.334delA | p.Ser112Valfs*14 | No | No | No |
| 3048 | FANCL | stop_gained | c.948T>G | p.Tyr316* | No | No | No |
| 251 | GEN1 | frameshift_variant | c.347_348delAG | p.Glu116Valfs*20 | No | No | No |
| 200226 | GEN1 | frameshift_variant | c.1933_1936delAAAG | p.Lys645Cysfs*29 | No | No | No |
| 1244 | MRE11 | stop_gained | c.1099C>T | p.Arg367* | No | Yes (1 missense mutation) | Yes |
| 2760 | MRE11 | stop_gained | c.1735C>T | p.Arg579* | No | No | No |
| 68 | MSH2 | frameshift_variant | c.2523dupA | p.Glu842Argfs*4 | No | Yes (1 nonsense mutation) | Yes |
| 3225 | MSH6 | frameshift_variant | c.3959_3962delCAAG | p.Ala1320Glufs*6 | No | Yes (1 frameshift insertion) | Yes |
| 92 | MUTYH<br>(monoallelic loss) | missense_variant | c.1187G>A | p.Gly396Asp | No | No | No |
| 213 | MUTYH<br>(monoallelic loss) | missense_variant | c.536A>G | p.Tyr179Cys | No | No | No |
| 280 | MUTYH<br>(monoallelic loss) | missense_variant | c.1187G>A | p.Gly396Asp | No | No | No |
| 353 | MUTYH<br>(monoallelic loss) | missense_variant | c.1187G>A | p.Gly396Asp | No | No | No |
| 442 | MUTYH<br>(monoallelic loss) | splice_region_variant | c.933+3A>C |  | Yes | No | Yes |
| 627 | MUTYH<br>(monoallelic loss) | missense_variant | c.1187G>A | p.Gly396Asp | No | No | No |
| 2365 | MUTYH<br>(monoallelic loss) | missense_variant | c.536A>G | p.Tyr179Cys | No | No | No |
| 2939 | MUTYH (monoallelic loss) | missense_variant | c.1187G>A | p.Gly396Asp | Yes | No | Yes |
| **3227** | *MUTYH*<br>(monoallelic loss) | missense_variant | c.1187G>A | p.Gly396Asp | No | No | No |
| **200193** | *MUTYH*<br>(monoallelic loss) | disruptive_inframe_deletion | c.1437_1439delGGA | p.Glu480del | No | No | No |
| **227039** | *MUTYH*<br>(monoallelic loss) | missense_variant | c.1187G>A | p.Gly396Asp | No | No | No |
| **587** | *PALB2* | frameshift_variant | c.509_510delGA | p.Arg170Ilefs*14 | No | No | No |
| **200127** | *PALB2* | stop_gained | c.2576C>A | p.Ser859* | No | No | No |
| **262114** | *PALB2* | splice_region_variant | c.49-2A>T |  | No | No | No |
| **352566** | *PMS2* | frameshift_variant | c.1831dupA | p.Ile611fs | Yes | No | Yes |
| **390** | *PMS2* | stop_gained | c.1687C>T | p.Arg563* | Yes | No | Yes |
| **101930** | *POLH* | stop_gained | c.328G>T | p.Glu110* | No | No | No |
| **2946** | *RECQL4* | frameshift_variant | c.871delG | p.Ala291Leufs*2 | No | No | No |
| **2957** | *RECQL4* | frameshift_variant | c.2492_2493delAT | p.His831Argfs*52 | No | No | No |
| **2768** | *SLX4* | stop_gained | c.2384C>G | p.Ser795* | No | No | No |
| **200019** | *TP53* | frameshift_variant | c.681delT | p.Asp228Thrfs*19 | Yes | Yes (1 splice site) | Yes |
| **200107** | *TP53* | missense_variant | c.743G>A | p.Arg248Gln | Yes | No | Yes |
| **4430** | *XPA* | frameshift_variant | c.103dupG | p.Ala35Glyfs*27 | No | No | No |
| **587** | *XRCC3* | frameshift_variant | c.606_607delGT | p.Arg204Glyfs*18 | No | No | No |

